# Genomic analyses in *Drosophila* do not support the classic allopatric model of speciation

**DOI:** 10.1101/2024.05.20.595063

**Authors:** Leeban H. Yusuf, Dominik R. Laetsch, Konrad Lohse, Michael G. Ritchie

## Abstract

The allopatric model of speciation has dominated our understanding of speciation biology and biogeography since the Modern Synthesis. It is uncontroversial because reproductive isolation may readily emerge as a by-product of evolutionary divergence during allopatry unopposed by gene flow. Recent genomic studies have found that gene flow between species is common, but whether allopatric speciation is common has rarely been systematically tested across a continuum of closely-related species. Here, we fit a range of demographic models of evolutionary divergence to whole-genome sequence data from 93 pairs of *Drosophila* species to infer speciation histories and levels of post-divergence gene flow. We find that speciation with gene flow is common, even between currently allopatric pairs of species. Estimates of historical gene flow are not predicted by current range overlap. Whilst evidence for secondary contact is generally limited, a few sympatric pairs showed strong support for a secondary contact model. Our analyses suggest that most speciation processes involve some long-term gene flow, perhaps due to repeated cycles of allopatry and contact, without requiring an extensive allopatric phase.

---

The idea that speciation processes can be classified into distinct geographic modes (allopatric, parapatric, and sympatric) emerged during the Modern Synthesis and continues to be a central theme of evolutionary research [48, 15, 47]. Allopatric speciation is thought to be common and is often considered a null model of speciation [15, 51]. Allopatric speciation appears uncontroversial both empirically given the widespread prevalence of ‘geographic’ varieties and species (which influenced Darwin and Mayr), and conceptually due to the ease with which the absence of gene flow facilitates the build-up of reproductive isolation (RI) due to genetic drift, natural, or sexual selection. Much of the debate over other modes of speciation revolves around the appearance of different forms of RI in the face of potentially homogenising gene flow. If and when species come into secondary contact after an allopatric phase, natural selection to reduce competition (but see [3]) or to prevent maladaptive hybridization (‘reinforcement’) may lead to complete isolation or the formation of stable hybrid zones [1].

Comparisons between currently sympatric and allopatric species pairs have provided apparently axiomatic evidence for reinforcement. In particular, analyses that contrast estimates of pre- and postzygotic RI between currently allopatric and sympatric species pairs of *Drosophila* demonstrated convincingly that mate discrimination evolves more rapidly between sympatric species [13, 14]. Alongside other comparative surveys [63, 64, 52, 23, 58], these patterns were interpreted as evidence that incompatibilities which arose during an allopatric phase lead to reinforcement after secondary contact [45]. However, classifying closely related species based on their current geographic ranges assumes that these in part reflect the geographical context during the key phases of their speciation history [39, 5, 29]. It has been acknowledged that allopatry and sympatry are ends of a spectrum [20, 44, 9], and an increasing number of genomic studies suggest that historical and contemporary gene flow is more prevalent than the architects of the modern synthesis had envisaged [56, 18, 21, 53, 17]. Additionally, gene flow may diminish gradually without the need for extended periods of allopatry during divergence [51]. It is therefore, unclear whether most speciation processes involve a strictly allopatric phase, and how common allopatric speciation versus speciation with gene flow is.

Here, we take advantage of the wealth of genomic data available for one of the best-studied groups of species, *Drosophila*. Population genetic analyses of *Drosophila* species pairs have spurred the development of inference methods that model the joint effects of incomplete lineage sorting and gene flow on local genealogies. These methods integrate over all possible genealogies sampled from short blocks of sequence across the genome, enabling maximum likelihood estimation of contrasting, nested models of species divergence and gene flow [31, 40, 33]. For small samples it is possible to calculate likelihoods analytically [61, 38], which allows efficient inference of simple demographic histories even froma single genome per species [61, 62]. While gene flow analyses in *Drosophila* have demonstrated its prevalence [56], we lack a systematic evaluation of the relationship between inferred historical demography and current range overlap in the genus. We analyse the pairwise distribution of sequence differences (which is a function of the distribution of pairwise coalescence times) in putatively neutral, short intronic blocks in 93 *Drosophila* species pairs to estimate the relative support for allopatric species divergence and speciation with gene flow. Sampling a single intronic block per gene justifies the assumption of statistical independence between blocks and allows us to calculate support for alternative demographic models of species divergence and parameter estimates in a likelihood framework [61, 62]. Our focal species pairs are a subset of the species pairs originally studied by Coyne and Orr [13, 14] and include currently allopatric and sympatric species spanning a range of genomic divergence (0.01 < *d*_xy_ < 0.05) (Supplementary table 2, and Supplementary Fig. 1 and 2). While our choice of taxon pairs was determined by the availability of whole genome sequence data, we note that our focal taxa cover a wide taxonomic and geographic range within the *Drosophila* genus [19]. Furthermore, the classic result of Coyne and Orr [13, 14] (faster evolution of pre-zygotic isolation for sympatric pairs) holds for this subset (Supplementary Fig. 1). In fact, both premating and postzygotic RI are higher in sympatric compared to allopatric pairs (two-sample Wilcoxon tests: W = 638.5, p = 0.004 and W = 94.5, p = 0.009), apattern already reported by [63, 46].

We fitted alternative demographic models of strict allopatric divergence versus divergence with gene flow and secondary contact [61, 62] (Fig. 1) to ask the following questions: (1) How much support for (historical or contemporary) gene flow is there and does this differ between currently sympatric or allopatric species pairs? (2) Do currently sympatric and allopatric pairs differ in their species divergence time and/or ancestral population size? (3) Finally, do sympatric pairs show greater support for secondary contact histories than allopatric pairs? We conclude that speciation involving a low rate of long-term gene flow is ubiquitous in *Drosophila*, and therefore that there is little support for allopatric speciation in this group.

**Fig. 1.**
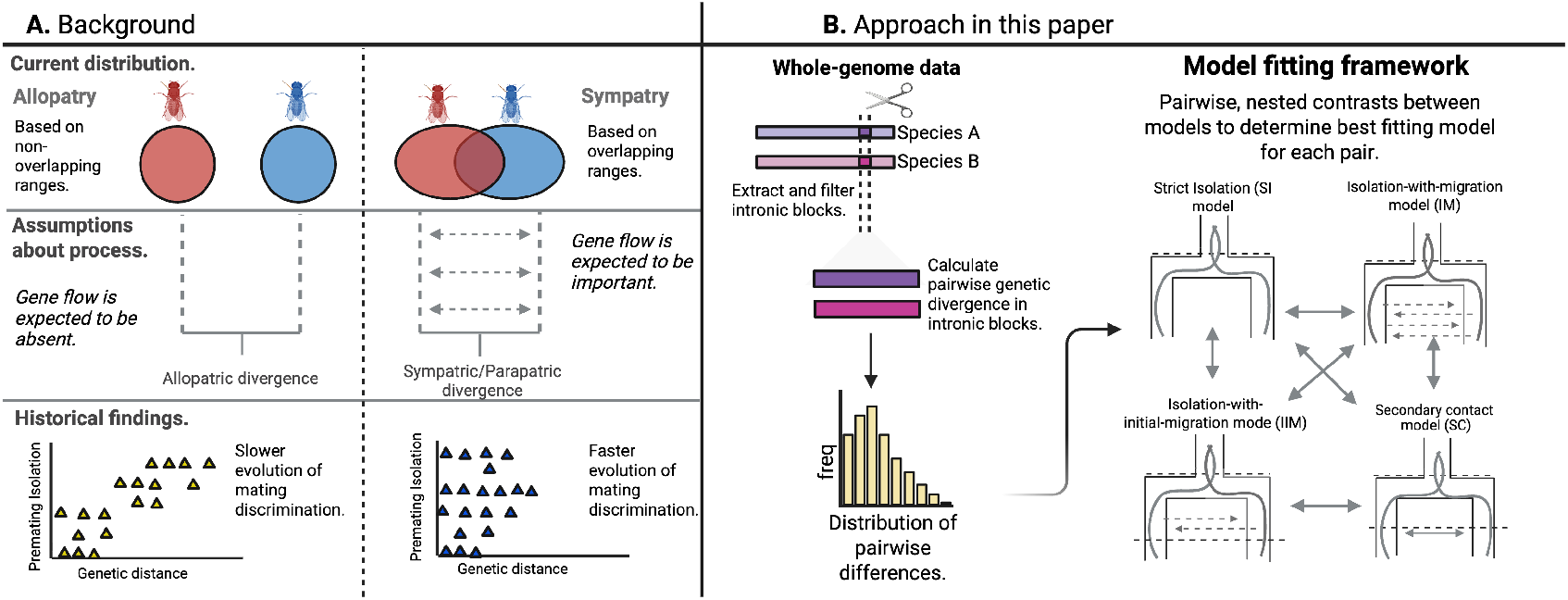
**A)** Background and **B)** our approach to understanding biogeographic modes of speciation. **A)** Previous comparative surveys on the evolution of reproductive isolation categorized species pairs by geography. This assumes that current range overlap is informative about the underlying speciation process, e.g. sympatric pairs are more likely to have exchanged migrants than allopatric taxa. **B)** Our approach uses whole-genome data to test to what extent levels of long-term gene flow differ between taxon pairs with and without current range overlap. We summarized genome-wide divergence in short, intronic blocks in terms of the distributions of pairwise differences. We fitted demographic models of speciation representing strictly allopatric speciation and different models of speciation with gene flow, to each pair and assessed their relative support in a likelihood framework.

## Results

### Speciation with gene flow is common across *Drosophila* pairs

To assess support for gene flow in each focal species pair of *Drosophila*, we compared four speciation models: (1) a strict isolation model (SI), where a pair of species diverge from an ancestral population of size *N*_*e*_ at some time *T*_0_ and remain in strict allopatry, (2) an isolation-with-migration model (IM), where divergence occurred with symmetric migration at a constant rate *M* = 2*N*_*e*_*m* per generation from *T*_0_ until the present and (3) a secondary contact model (SC) where divergence occurred without gene flow initially, but where the pair experiences an instantaneous burst of bidirectional gene flow that transfers a fraction *f* of lineages at time *T*_1_ (see SI Appendix section 1.4 for details). The latter may reflect a ‘reinforcement’ scenario where gene flow only occurs transiently following secondary contact. We restrict analyses to these relatively simple models both because previous comparative surveys of speciation demography have considered the same scenarios [53] and because more complex models are unlikely to be identifiable from the pairwise distribution of differences. In particular, we also considered an isolation-with-initial-migration model (IIM) where gene flow occurred at the onset of divergence but stopped at time *T*_1_. However, since only one taxon pair (*D. lummei* - *D. littoralis*) had significantly support for IIM, we concluded that this scenario is either not identifiable or not sufficiently supported by the data and so restricted comparison of demographic histories to SI, IM and SC models. Note also that our gene flow estimates are effective parameters which incorporate any selection against early generation hybrids (see Discussion).

Assigning a best-fitting but minimally complex model to every species pair reveals overwhelming support for speciation histories that involve gene flow: out of the 93 *Drosophila* pairs, only 12 pairs best fit an SI model, i.e. neither IM nor SC give a significant improvement in model fit (Fig 2A), and four of these are sympatric. For the majority of species pairs (76) both IM and SC scenarios fit significantly better than a SI history and relative support for either model is highly correlated (Fig. 2A). Given that the SC model is more complex than the IM model (a total of four instead of three parameters), we only accepted it as the best and most parsimonious history when the relative support for it exceeded a critical value of 2Δ ln *L* > 3.841. We also used this threshold (which is equivalent to assuming a *χ*^2^ distribution with 1 d.f.) to compare nested models that differ by one parameter (IM vs SI and IIM vs IM). Given these model selection criteria, we find that an SC history is the best supported model for eight pairs and accept the IM model as the best-fitting history for 72 pairs (Fig. 2A).

**Fig. 2.**
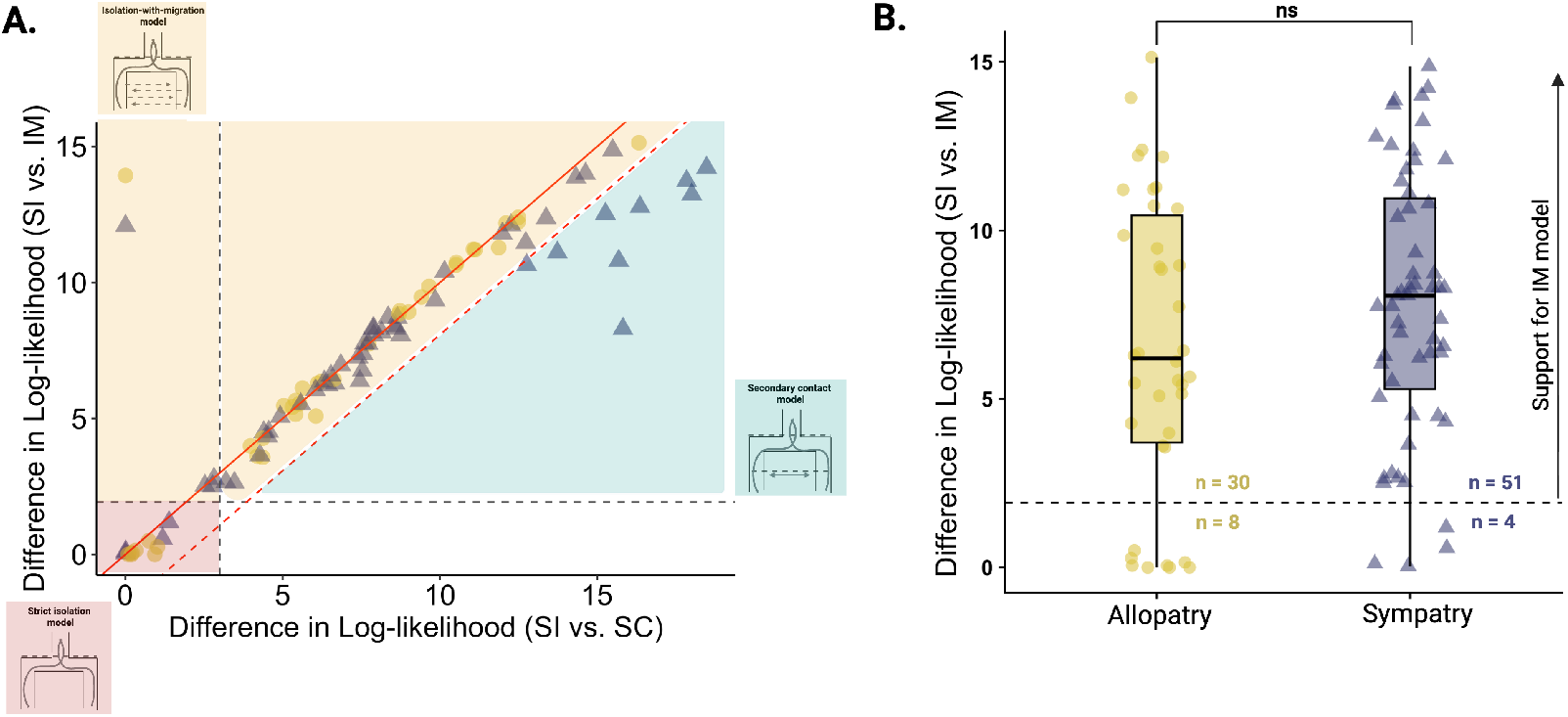
Evidence for gene flow across allopatric and sympatric *Drosophila* pairs. This plot summarises which pairs best-fit the SC model (right bottom, teal) IM (top right, yellow) and SI (bottom corner, red). The vast majority of pairs fall into regions supporting gene flow. **A**. The difference in log-likelihood between strict isolation (SI) and isolation-with-migration (IM) histories against the difference in log-likelihood between strict isolation (SI) and secondary contact (SC) histories. Black-dashed lines are the critical value thresholds for the SI v.s. IM (horizontal line at 2Δ ln *L* > 3.84, 1 d.f.) and SI v.s. SC (vertical line at 2Δ ln *L* > 5.99, 2 d.f.). The red solid line is the line of equality (y=x), showing correlated support for gene flow models regardless of whether gene flow is continuous or an instantaneous event. Points to the right of the dashed red line represent pairs where support for the SC model exceeds (a critical value of 2Δ ln *L* > 3.84) the support for an IM model, with the exception of a single pair which best fits an IIM history better than both the IM and SC models. Two pairs fit the IM model considerably better than the SC model (top left). Shaded areas denote best-fitting models in our comparative framework. **B**. The difference in relative support for an IM history, against a SI history, for currently sympatric and allopatric species pairs. The vertical dashed line indicates the critical value (p < 0.05). The numbers of species pairs that fit or do not fit an IM model significantly better than an SI model are shown on either side of the vertical line.

### No difference in support for historic gene flow between sympatric and allopatric *Drosophila* pairs

If current range is indicative of the mode of speciation, we would expect allopatric pairs to best fit a history of strict isolation (SI) with no significant improvement in fit when including gene flow. In contrast, we may expect currently sympatric pairs to fit an isolation-with-migration (IM) model better than an SI model. Alternatively, if sympatry is recent (and associated with secondary gene flow), we may expect sympatric pairs to be associated with histories of secondary contact (SC) (‘recent sympatry’ hypothesis).

We find no difference in relative support (as measured byΔ ln *L*) for an IM model over an SI model between allopatric and sympatric pairs (two-sample Wilcoxon test: W = 867, p= 0.165) (Fig. 2A). However, we do find that the 12 pairs that do not show support for any gene flow scenario, i.e. that best fit an SI history, include more allopatric than sympatric pairs (one-tailed Fisher exact test: p = 0.064). Interestingly, these SI pairs almost entirely belong to the *Drosophila* nasuta group, including the species pair *D. albomicans* and *D. nasuta*, which in previous analyses also did not show evidence for post-divergence gene flow [4]. All eight pairs that best fit a secondary contact (SC) model are currently sympatric species, consistent with our ‘recent-sympatry’ hypothesis (one-tailed Fisher exact test: p = 0.019) (Fig. 2A)

### Current range overlap poorly predicts the demography of speciation

To test whether historical gene flow differs between allopatric and sympatric pairs, we converted estimates of the scaled effective rate of gene flow (i.e. the number of migrants per generation *M*) under the IM model, into a per lineage probability of gene flow: e.g. given an estimated duration of gene flow between species *T*, the probability that the ancestry of an individual haplotype at a random position in the genome is affected by migration is 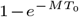. This conversion allows for a direct comparison between estimates of continuous (IM) and discrete (SC) gene flow. We find a lower long-term probability of gene flow for allopatric pairs (mean probability of gene flow: 66%) compared to sympatric pairs (mean probability of gene flow: 74%). When assuming the best-fitting gene flow modelfor each pair, this difference is not significant (two-sample Wilcoxon test: W = 934, p = 0.387)(Fig. 3A). Hence, there is evidence of less gene flow between allopatric species, though the extent is still high. Note that the IM model, although accepted as the best supported model for 72 taxon pairs yielded nonsensical parameter estimates in 22 pairs, i.e. we obtained arbitrarily high estimates of either gene flow (*M*) or divergence time *T*_0_. This suggests either poor model fit and/or parameter non-identifiability (see Discussion) and we have removed these estimates from subsequent analyses. Excluding the 22 pairs with non-identifiable migration parameters did not change the lack of difference in long-term gene flow probability between sympatric and allopatric pairs (two-sample Wilcoxon test: W= 603, p = 0.847).

**Fig. 3.**
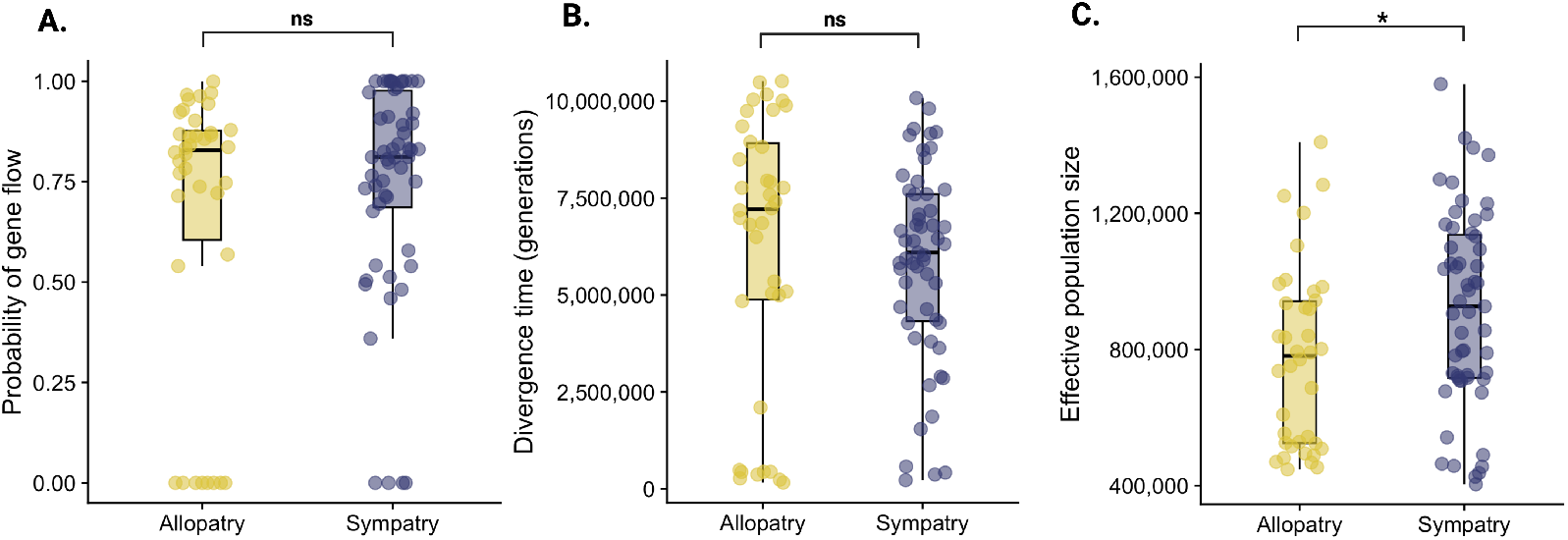
Sympatric and allopatric pairs show similar divergence times, effective population size and levels of long-term gene flow. Current range overlap does not reflect inferred historical demography. **A**. We find no difference in long-term rates of gene flow between allopatric and sympatric pairs. Gene flow was calculated for each pair using parameter estimates from the best-fitting model. **B**. and **C**. We find no difference in scaled divergence time, but we find a difference in effective population size (the best-fitting model for each pair) between allopatric and sympatric pairs. ‘ns’ indicates no significant difference and ‘*’ indicates p < 0.05 via Mann-Whitney U test.

Additionally, we find that currently allopatric and sympatric pairs do not differ in the onset of divergence (measured in generations), but do differ in ancestral effective population size (Fig. 3B and C). While sympatric pairs are younger on average than allopatric pairs, this difference is not significant (two-sample Wilcoxon test: W = 1219, p = 0.175) (Fig. 3B). However, sympatric pairs have larger ancestral effective population sizes than allopatric pairs (two-sample Wilcoxon test: W = 768, p = 0.03). Note that this difference is not significant when we exclude the 22 species pairs with non-identifiable parameters (two-sample Wilcoxon test: W = 458, p= 0.06).

### Secondary contact pairs do not have greater pre-mating isolation

Reinforcement is frequently invoked to explain enhanced mating discrimination in sympatry relative to allopatry [49, 15]. We asked whether the eight pairs that fit a secondary contact (SC model) substantially better than an isolation-with-migration (IM) model (2Δ ln *L* > 3.841) show greater premating isolation, than pairs for which IM is the best fitting model (Δ ln *L* > 0; n = 72). We find no significant difference in premating isolation for SC pairs (mean pre-mating isolation:: 0.90) compared to pairs that best fit an IM model (mean pre-mating isolation: 0.88) (two-sample Wilcoxon test: W = 294.5, p = 0.816).

## Discussion

Allopatric speciation is considered the most common mode of speciation because the evolution of RI is unhindered once gene flow has ceased completely [15]. Analysing genomic data for 93 *Drosophila* species pairs in a hierarchical modelling framework, we find that most pairs show evidence for post-divergence gene flow. Perhaps surprisingly, while we find some evidence for increased gene flow between currently sympatric pairs, our main finding is that some level of post-divergence gene flow is common, even between currently allopatric pairs. While sympatric pairs show overall greater support for secondary contact histories and very recent gene flow as expected, there is little or no difference in statistical support for histories involving gene flow between pairs that are currently sympatric or allopatric.

These results have implications for interpreting classic comparative surveys of speciation in *Drosophila* and other taxa. Mating discrimination has been shown to evolve more rapidly in currently sympatric *Drosophila* species pairs relative to allopatric pairs, and this observation has been interpreted as support for reinforcement [15, 45, 54, 50]. The fact that we find significantly greater support for secondary contact histories for sympatric compared to allopatric species pairs is compatible with a role of reinforcement in speciation and shows that our analyses are extracting genuine signals of gene flow. In order to understand which aspects of the data allow us to distinguish between SC and IM models, it is helpful to inspect their absolute fit to the observed distribution of pairwise differences: taxon pairs that best fit an SC history show a characteristic excess of monomorphic blocks (*S* = 0) in the data (Supplementary Figs 13-15). While this high frequency of monomorphic blocks can be explained by a recent burst of admixture upon secondary contact, it is incompatible with histories of continuous gene flow (i.e. the IM model).

### Limits and robustness of demographic inference

Our core finding that overall levels of historic gene flow differ little between currently sympatric and allopatric *Drosophila* pairs is perhaps surprising. It is important to stress that our coalescence-based inference of demographic history is based on minimal sampling of a single haplotype per species and is therefore limited to a small set of simplistic models. The fact that IM and SC models, which assume symmetric migration and ignore heterogeneity in *N*_*e*_, already provide a good absolute fit to the observed *S* distributions suggests that more realistic models that include multiple phases of gene flow or account for asymmetry are not identifiable. Future population genomic analyses that include data on intraspecific diversity will undoubtedly be able to fit more realistic demographic models of species divergence.

There are, however, fundamental limits to demographic inference that do not depend on the sample sizes and data summaries used: estimates of gene flow and ancestral *N*_*e*_ between taxon pairs are potentially confounded by ghost admixture into one or both focal taxa from a third taxon [6]. Furthermore, periods of high gene flow (or low *N*_*e*_) erase the genomic footprints of older demographic events and inference is limited to long-term rates of gene flow that are sufficiently low (*M* < 10). Importantly, this means that periods of gene flow that are high enough for reinforcement selection to act may be indistinguishable from panmixia.

Given the simplicity of our data summary, an important question is to what extent our finding of pervasive historic gene flow in *Drosophila* hinges on the ability to accurately estimate the frequency of monomorphic blocks. This may be difficult for two reasons: firstly, intronic sequence may be incompletely annotated in some taxa and including conserved regulatory sequence in introns will inflate the frequency of monomorphic blocks. Secondly, many of the sequence data we analyse are from isofemale lines with a history of lab culture and it is perhaps feasible that in some pairs recent admixture may have occurred as a result of lab contamination. To assess the dependence of our results on the frequency of monomorphic blocks, we estimated the fit of SI, IM and SC models for all pairs when monomorphic blocks are excluded. This involves conditioning on only observing blocks that differ between species in a pair by maximizing the conditioned loglikelihood ln 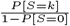. We find that while conditioning on*S* > 0 – unsurprisingly – decreases support for IM and SC histories (relative to SI) overall (see Supplementary Fig 13-15; SI model: 15 (allopatric) and 24 (sympatric) pairs; IM model:23 (allopatric) and 31 (sympatric) pairs; SC model: 0; IIM model: 0), our main qualitative result of a signature of gene flow in most taxa (n = 54) without any systematic differencebetween sympatric and allopatric pairs is unaffected (Δ ln *L* between SI v.s. IM models; two-sample Wilcoxon test: W = 1242, p = 0.124).Finally, a power analysis based on data simulated under plausible parameters for *Drosophila* shows that the overall greater support for speciation histories involving a low level of gene flow is unlikely to reflect biases that arise from the fact that our multilocus inference framework ignores recombination within blocks (see simulation results in Supplementary section1.5 and Supplementary Fig. 8). Additionally, when we use a simple phylogenetic correction to account for the non-independence of some species pairs, we still find consistent evidence of gene flow across nodes in the *Drosophila* phylogeny [2](Supplementary section 1.6; Supplementary Fig. 11 and 12). Thus, there is a still pervasive signal of post-divergence gene flow in *Drosophila* pairs, even between currently allopatric pairs, and across all of our sensitivity analyses.

### Speciation with gene flow is the rule not the exception

Our results strongly imply that speciation does not require an extended allopatric phase to allow the build-up of RI. Instead, a low level of gene flow at all stages of speciation appears to be relatively common in *Drosophila*. Numerous other studies have inferred gene flow between ‘good’ species without explicitly testing if this is greater in sympatric species [41, 65, 56, 53]. It appears that speciation in the face of gene flow is common and the traditional classification into distinct geographic modes of speciation, with allopatry taken as the default, is outdated [44, 9, 20, 42]. The importance of ‘strict isolation’ to the Biological Species Concept has diminished greatly over the last few decades, partly as a result of genomic analyses of introgression. Moreover, it is generally accepted that speciation is a continuous process [55, 43] and that complete RI is not necessary between species. Instead, our results suggest that speciation more often involves extended periods of genetic exchange and incomplete RI rather than an abrupt and complete cessation of gene flow. Of course, this widens the ‘grey zone’ where species barriers remain permeable long after species divergence has become irreversible [43, 7]. Genomic analyses of speciation with gene flow clearly demonstrate that levels of gene flow can vary widely across the genome [34, 22]. The extent to which this variation reflects clustered genetic architectures for barriers to gene flow that have arisen as a consequence of speciation with gene flow or simply pre-existing variation in recombination, perhaps owing to structural rearrangements such as inversions, is an open question. Nevertheless, it is clear that species barriers can build up and reach tipping points in the face of on-going gene flow without any extended period of strict allopatry.

However, it is important to emphasise that our suggestion that a strict allopatric model of speciation is probably uncommon in *Drosophila* does not suggest that allopatry is unimportant or plays no role. Studies of phylogeography, ancient DNA and niche modelling recognise that species ranges may be dynamic over timescales that are short relative to speciation processes. Within the limits of our modeling approach we have demonstrated that some amount of long term gene flow during species divergence is almost ubiquitous. However, estimating the timing and duration of individual episodes of gene flow is arguably a much more difficult task. The Pleistocene climate history has been dominated by cycles of ice ages and warmer interglacials. Thus, species both in the temperate zones [30, 28] and the tropics [26, 11] have undergone repeated periods of allopatry and secondary contact and gene flow. Such complex histories would be challenging to model explicitly (and are impossible to infer from the distribution of pairwise differences). By contrast, the IM model assumes a single long term (over the scale of *N*_*e*_ generations) rate of gene flow. This parameter therefore reflects a long term average of genetic exchange over many Pleistocene cycles of range shifts.

Another important consideration is how applicable our results are to other taxa. *Drosophila* is highly vagile, flighted and long distance dispersal has been demonstrated in some species [12]. However, many *Drosophila* species have specialised niches, so successful dispersal requires either movement of host plants, commensalism or switches to novel hosts. One may expect larger, less vagile, organisms to show less dynamic rangechanges. However, phylogeographic patterns of diversity that imply rapid range shift in response to glacial cycles have been recognised in vastly diverse taxa, including plants and larger vertebrates [30]. We therefore see no reason to suppose that our result that current range overlap is essentially uncorrelated with historic gene flow during species divergence is limited to *Drosophila*.

We stress that demographic analyses of larger genomic datasets (preferably from wild-caught samples and known areas of range overlap) that make use of intraspecific variation will allow to fit more realistic models of divergence and may help to answer new questions. For example, rather than assuming that gene flow during species divergence is symmetric – as we have done here – it would be fascinating to test which traits and population genetic processes correlate with the direction of gene flow. There is clearly also a need for similar comparative analyses of speciation histories for taxa that have richer geographic range and life history information than *Drosophila* to contextualise the evolution of RI and the extent, timing and direction of gene flow at different points along the speciation continuum.

## Methods

### Data sampling and quality control

We collated data on genetic distance, reproductive isolation (RI) and range overlap for 93 pairs of Drosophila species from published datasets [63, 64, 13, 14] (see Supplementary table 5 for RI data). These pairs are relatively phylogenetically broad, as we include species pairs from the melanogaster, repleta, virilis, immigrans, obscura, and willistoni groups. According to a recent *Drosophila* phylogenies [32, 19], these lineages are well distributed across the genus. We augmented these with analyses of demographic history for each species pair using genomic data to test key assumptions about the relationship between current range overlap and speciation history. To do this, we obtained one genome assembly (based mostly on long read data) per species pair and whole-genome sequencing (WGS) data (Illumina, short read) for each species from publicly available datasets via NCBI (Supplementary table 3). We only included WGS data with similar coverage, removed pool-sequenced or experimentally manipulated samples, and prioritised data obtained from wild-caught individuals (rather than lab lines). For WGS datasets that met these criteria, we randomly assigned one resequencing dataset to represent each species. Full filtering details can be found in SI Appendix (section 1.1). Details on genome assemblies and WGS datasets sampled can be found in Supplementary table 3 and 4 in the SI Appendix.

### Genome annotation, mapping and variant calling

We annotated each genome separately using BRAKER2 (v2.1.6) and *D. melanogaster* protein sequences as evidence [8]. Raw reads for each WGS dataset were trimmed using fastp [10], aligned using bwa-mem2 [36, 59], and finally sorted and filtered using sambamba [57] and picard. Both WGS datasets of each species pair were mapped to the annotated genome (belonging to one of the two species) in the pair. Variants were called using freebayes and filtered for missing genotypes, read mapping bias and depth using bcftools and gIMble[24, 35, 34, 16]. For full details on annotation, mapping and filtering see SI Appendix (section 1.2).

### Sampling intronic blocks

We opted to use intronic sequences for demographic analyses since introns can be more reliably annotated than intergenic regions but show much less functional constraint than protein-coding genes. Functional constraint across introns can be variable, so we implemented a range of filtering strategies to keep intronic segments most likely to be selectively neutral[27, 25]. To ensure we minimised linkage between intronic regions, we only sampled one intronic block per gene. Intronic blocks of a fixed, species-specific length were sampled from callable variants after filtering. This means that blocks could span intronic sequence (within a single intron) that were excluded because quality or coverage filters. Our filtering strategy is described in full detail and can be found in the Supplementary Material (section 1.3 and Supplementary Fig. 9). As a sanity check on our filtering strategy we compared *d*_XY_ in the filtered intron dataset to *d*_XY_ calculated across all sites in the genome (Supplementary Fig. 10). This confirms that selective constraint on the intronic sequences included in our analyses is low compared to the genome-wide average.

Extracting genomic DNA for a single *Drosophila* sample for sequencing often requires pooling multiple individuals from an isofemale line. Even when a single individual can be sequenced, the heterozygosity in the resulting WGS data reflects both the effective size (*N*_*e*_) at the species level and the history of lab culture. We therefore restrict our analysis to the most minimal sampling scheme of a single haplotype per species and base inference on the distribution of pairwise differences (*S*) between species. The *S* distribution in short intronic block is a vector of counts 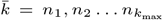). We assumed thatthe heterozygous sites in each block can be randomly assigned to haplotypes using a simple binomial sampling procedure (implemented in `Mathematica` v12.3) that considers all ways of phasing heterozygous sites: e.g. a block containing two fixed differences between species and a single heterozygous site contributes probabilities 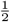 each to *S* = 2 and 3. Likewise, a block with two fixed differences and two heterozygous sitescontributes probabilities 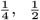 and 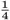 to *S* = 2, 3 and 4respectively. To account for differences in the number of blocks between species pairs, we normalised *S* distributions by 500*/n*_*i*_ (where *n*_*i*_ is the number of blocks in pair *i*).

### Modelling speciation histories across *Drosophila*

To understand demographic histories and the degree of gene flow for species pairs, we fitted a range of models of demographic history to the distribution of pairwise differences in short intronic blocks between species: (a) a strict isolation (SI) model (most consistent with strict allopatric speciation) characterised as an instantaneous split of an ancestral population at *T*_0_ without gene flow, (b) isolation with migration (IM), where an ancestral population diverges with symmetric migration (M = 4*N*_*e*_m migrants per generation, where *N*_*e*_ is the ancestral population) between the time of divergence and the present, (c) an isolation with initial migration model (IIM), where an ancestral population diverges with an initial period of symmetric migration and gene flow ceases at *T*_1_ and (d) a secondary contact (SC) model, where an ancestral population diverges in allopatry, and an instantaneous recent, pulse of gene flow, i.e. a total proportion (f) of the population is introgressed at time *T*_1_. This secondary contact model differs from the other SC models in the literature in that gene flow is modeled as a short bidirectional pulse rather than continuous migration from *T*_1_ to the present [53], reflecting the expectation that reinforcement rapidly halts gene flow during secondary contact. In all cases, we assumed a single *N*_*e*_ parameter that is shared between the ancestral population and the daughter species. See Supplementary Figure 7 for visualisations of each model.

Analytic solutions derived in [61, 38, 62] allow efficient maximum likelihood estimation of parameters under the IM and IIM model from the *S* distribution. We use the expression for the probability of seeing *k* differences between a pair of sequences sampled from different populations under the IM model and IIM model *P* [*S* = *k*] [61, eq. 24] and [62, eq. 29]. We assumed a single *N*_*e*_ parameter *θ* = 4*N*_*e*_*µ* for the ancestor and the two daughter species and a fixed mutation rate across all blocks. The likelihood expression for the secondary contact model was adapted from [37]. For each model and species pair we maximized the log likelihood across blocks:

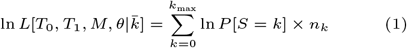

This likelihood calculation was implemented in Mathematica; we used the function FindMaximum to maximise eq. 1 and obtained maximum likelihood estimates (MLE) of parameters under each model and for each species pair. Given that the SI, IM and IIM models are nested, we used likelihood ratio tests (assuming the 2Δ ln *L* follows a *χ*^2^ distribution) to determine relative model support for each species pair. Since the IIM and SC models have the same number of parameters, we compared them simply in terms of relative log-likelihood. A detailed description of our rationale for evaluation of model support can be found in Supplementary Material (section 1.4.1 and 1.4.2).

To scale parameter estimates, we used the spontaneous mutation rate estimate of *µ* = 3.32 × 10^−9^ per base and generation for *D. melanogaster* and additionally included the95% confidence intervals (2.52 × 10^−9^ - 4.30 × 10^−9^) of this estimate (26).

### Simulations

A key assumption of our inference framework is that it is possible to sample neutrally evolving variation in short block of sequence within which recombination can be ignored. Violations of this assumption can lead to biases in parameter estimates and could result in erroneous support for histories of gene flow[60]. To rule out this possibility, we simulated *S* distributions for each pair in our dataset using the maximum likelihood parameter estimates under a SI model and realistic rates of recombination (*r* = 1.03 cM/Mb) and mutation 3.32 ×10^−9^ (38). Comparing Δ ln *L* between SI and IM histories for simulated datasets suggest a maximum false positive rate of25 %, far below the proportion of species pairs that support a history of gene flow (87 %). Moreover, we find that the IM model has considerably higher relative support in the real data compared to data simulated under a null model with recombination but no gene flow. Full details can be found in SI Appendix (section 1.5).

## Acknowledgments

We thank the reviewer and the associate editor for their feedback. We thank Sam Ebdon for bioinformatic advice. Additionally, we thank Roger K. Butlin and Jim Mallet for helpful feedback that improved the manuscript, and Cher Chow for assistance with figures. LY was supported by a University of St Andrews studentship. KL was supported by a fellowship from the Natural Environment Research Council (NERC, NE/L011522/1) and a European Research Council starting grant (ModelGenomLand 757648) which also supported DRL. MGR was supported by a grant from the Natural Environment Research Council (NE/V001566/1).

## Author contributions

Conceptualisation: LHY, KL, MGR. Analysis: LHY, DRL, KL. Writing (original draft): LHY, KL, MGR. Writing (review and editing): LHY, KL, MGR.

The authors declare that they have no competing interests.

