## Supplementary Text/Material for "Genomic analyses in *Drosophila* do not support the classic allopatric model of speciation"

**This PDF file includes:**

Supporting text
Figures S1 to S15
Tables S1 to S6
SI References

### Supplementary Text

#### *1.1 Data sampling and quality control.*

To infer the demographic histories of species pairs of *Drosophila*, we collated data on reproductive isolation (RI), range overlap and genetic distance for *Drosophila* species pairs originally collated by Coyne and Orr (*1*, *2*), and expanded by Yukilevich (*3*, *4*). This represents the largest collection of RI measurements in *Drosophila*. As in (*1*, *2*), and other comparative surveys of *Drosophila* (*3*–*7*), we refer to "species pairs" as closely related but not necessarily independent pairs of species (i.e. our dataset consisted of 93 species pairs involving 58 species). Our goal was to infer speciation histories for as many species pairs found in (*1*, *2*) as possible to ensure results were comparable with previous comparative analyses. Briefly, Coyne and Orr (*1*, *2*) calculated premating isolation for each species pair as the percentage of successful copulations for homotypic and heterotypic matings during no-choice, single-choice, or multiple-choice mating experiments. Postzygotic isolation was calculated as the percentage of F1 hybrid males and females that were sterile or inviable. For each reciprocal mating, measures ranged from 0 (F1 hybrids are viable and fertile) to 1 (all hybrids produced are either sterile or inviable). The overall postzygotic isolation index is the average of both reciprocal mating interactions. For detailed descriptions of RI and range measures see references (*3*, *4*).

We sought to combine data on RI and range overlap with inferences of the demographic history of species divergence based on whole genome sequence (WGS) data. For each focal species pair, our data consisted of a single reference genome assembly as well as additional short read WGS data for a single sample of each species. Both were obtained from publicly available datasets via NCBI. 22 of the reference assemblies were based on long read data that were generated as part of large-scale efforts to produce highly contiguous *Drosophila* genome assemblies (*8*) (see Supplementary Table 1-3); the remaining 8 assemblies were based on Illumina data. For species pairs for which multiple genome assemblies were available, we assessed genome quality (using BUSCO v5.1.2 single-copy completeness) and selected the genome with the highest BUSCO score (>90% single-copy completeness) (*9*) as the reference for each pair.

To standardize quality of WGS datasets, we filtered the possible sequencing datasets for each species (see Supplementary Table 1 for a list of samples used): specifically, we restricted our analyses to high coverage datasets (i.e. total number of bases between 1 and 50 Gb) and removed pool-sequenced samples or samples that had been experimentally manipulated before sequencing. We selected data from wild-caught individuals where possible. For each focal species, we randomly sampled a single WGS dataset using a custom R script from the list of publicly-available samples. The randomly sampled WGS data for each pair were downloaded using the sratools functions pre-fetch v2.11.0 and fasta-dump v2.11.0.

#### *1.2 Genome annotation, mapping and variant calling.*

Reference genomes were annotated in the same way to minimize differences in gene prediction. Annotation was performed for all genome assemblies using BRAKER2 v2.1.6 (*10*). We used only *D. melanogaster* (Flybase v.6.39) proteins as evidence for all annotations. Subsequently, we extracted intronic regions from each genome annotation using BEDOPS v2.4.39 and bedtools v2.30.0 (*11*, *12*).

All reads were trimmed using fastp v.0.20.1. The reference genome assembly for each pair was indexed (Supplementary Table 3) (*13*, *14*). Species-specific paired-end or single-end read WGS data were then mapped to pair-specific genome assemblies using bwa-mem2 (v. 2.2.1) for species with paired end reads and bwa-mem (v. 0.7.17) for species with single end reads (*13*, *14*). For 9 species pairs a representative genome for mapping and variant calling did not exist for either species. For these pairs we used a reference genome from a closely related outgroup species. Reads were sorted, duplicates mapped and read groups added using sambamba v. 0.8.1 and picard v2.26.3(*15*, *16*). Trimming and mapping were implemented in the workflow engine

Snakemake v. 6.5.1 (*17*). Variants were called for reads in each pair using Freebayes v1.3.5 and subsequently filtered for missing genotypes, read mapping bias (RPL<1 | RPR<1 | SAF<1 | SAR<1), and a minimum read depth of 2, using bcftools v1.11 and gIMble v.0.6.1 (*18–20*). We refer to pair-specific datasets which have been filtered as above as 'callable and filtered' datasets. The workflow for the filtering process is summarized in Supplementary Figure 9.

#### *1.3 Sampling intronic blocks.*

To generate comparable datasets for demographic modelling across all pairs, we sampled intronic blocks from pair-specific, callable and filtered datasets. Our rationale for using intronic blocks to model speciation history (Supplementary Figure 9), was to maximise information contained in a single genome per species: our inference scheme summarizes variation in short sequence blocks that are assumed to be non-recombining and to evolve neutrally. Thus, coding regions are not suitable for such block-wise inference given their functional constraint. Similarly, although intergenic regions show, on average, similar levels of sequence conservation to introns in *Drosophila*, distinguishing conserved from selectively-neutral intergenic regions is challenging and relies heavily on accurate annotation (*21–23*). In contrast, annotation of genic regions and introns is comparatively easier and more reliable across pairs. Since functional constraint varies across introns in *Drosophila*, we applied additional filters to the filtered callable datasets (*21, 23*). Specifically, first introns were removed because they evolve more slowly than non-first introns (*21*), likely owing to their higher abundance of regulatory elements. We also removed the first and last 10 bases from all introns in our analyses using bedtools v2.30.0, and a custom python script.

We intersected filtered intronic sequence with callable positions using bedtools intersect v2.30.0. To minimise the effect of linkage between blocks on our analysis, each gene could only contribute a single intronic block to each pair-specific dataset. In practice, however, for intronic sequence to be useable it also had to satisfy the filtering criteria detailed in subsection 1.2 and be sufficiently long; this meant that not all genes contributed an intronic block to the dataset for each pair. Intronic blocks for every species pair were also manually checked to confirm that only a single intronic block per gene was sampled. To confirm that our datasets of filtered intronic blocks were more selectively neutral than the background, we compared absolute genetic divergence $d_{XY}$ calculated across filtered introns to $d_{XY}$ across all callable sites in the genome. As expected, given the near absence of functional constraint in filtered introns, $d_{XY}$ in filtered introns was higher than genome wide $d_{XY}$ in all pairs (Supplementary Figure 10).

For each species pair, short blocks were partitioned using gIMble v.0.6.1. For every pair, the block length was chosen to give an average of three pairwise differences (pair-specific block length = $3/d_{XY}$) between species per block. Species pairs with fewer than 250 blocks and pairs for which the short intronic blocks had higher within-species heterozygosity compared to fixed differences between species were removed from the analysis. These two filters removed 9 species pairs from the analysis. For the remaining 93 pairs, block sizes ranged from 60 – 341 bases, and the number of blocks ranged from 367 – 6390 (Supplementary Table 2). The final datasets consisting of pair-specific, filtered, short intronic blocks were converted into blockwise site-frequency spectra (bSFS), containing blockwise counts of all possible mutational configurations, using gIMble (v.0.6.1) which were further summarised into S distributions by assigning heterozygous sites to haplotypes at random assuming binomial sampling (see Methods). To account for differences in the number of blocks between species pairs, we normalised each S distributions by $500/n_i$ (where $n_i$ is the number of blocks in pair i). This is analogous to taking an average across all subsamples of 500 blocks for each taxon pair.

#### *1.4 Modelling gene flow and divergence.*

To understand demographic histories and the degree of gene flow for species pairs, we fitted a range of models of demographic history to the S distribution of each species pair : (a) **a strict isolation (SI) model** (most consistent with *strict* allopatric speciation), i.e. an instantaneous split of an ancestral population of effective size $N_e$ at time $T_0$ without gene flow, (b) **isolation with**

**migration (IM)**, where an ancestral population diverges with symmetric migration ($M = 4N_e m$ migrants per generation) between the time of divergence and the present, (c) an **isolation with initial migration model (IIM)**, where an ancestral population diverges with an initial period of symmetric migration and gene flow ceases at $T_1$ and (d) **a secondary contact (SC) model**, where an ancestral population diverges in allopatry, but experiences an instantaneous recent, pulse of gene flow, i.e. a total proportion (*f*) of the population is introgressed at time $T_1$. In all cases, we assumed a single $N_e$ parameter that is shared between the ancestral population and the daughter species. See Supplementary Figure 7 for visualisations of each model.

#### *1.4.1 Evaluating difference in support for gene flow between pairs based on current range overlap.*

Statistical analyses were performed in R using the package stats v.4.0.2, and in *Mathematica* (*29*). We hypothesised that if current range overlap is predicted by the demographic history of speciation, sympatric pairs should (1) fit an IM model better than an SI model more often than allopatric pairs, and (2) show greater support for an IM model over an SI model compared to allopatric pairs. We tested these two predictions by calculating the difference in log-likelihood (ΔlnCL) between the IM model and SI model. Pairs showed support for an IM model over an SI model if ΔlnCL was larger than the critical threshold ($p < 0.05$). We compared the number of allopatric and sympatric pairs fitting an IM model over an SI model using Fisher's exact test. To evaluate whether there was a difference in average support (ΔlnCL) for an IM model over an SI model between allopatric and sympatric pairs, we used *Mann-Whitney U* tests. This framework was used to test the difference in model support between allopatric and sympatric pairs, and difference in the proportion of pairs fitting a given model over another.

We used *Mann-Whitney U* tests to assess differences in the estimates of the scaled migration rate (M) between allopatric and sympatric pairs across models with gene flow (IM, IIM, SC). Additionally, we computed the probability that an individual lineage is involved in gene flow as $1 – e^{-\tau 0 * M}$ under the IM and IIM model. This allows comparisons of M estimates obtained under the IM and IIM model with the admixture fraction (*f*) computed in our SC model, which is already a per lineage probability. Similarly, we asked whether currently allopatric and sympatric pairs differed in the average probability of migration and/or the scaled divergence times and effective population sizes using *Mann-Whitney U* tests.

#### *1.4.2 Assigning a best-fitting model to each species pair.*

To assign the best-fitting model to each species pair, we compared model support pairwise between competing models. Critical values for the difference in support (ΔlnL) between nested models (at a significance level of 0.05) were determined assuming a $\chi^2$ distribution, which is conservative. For example, the SI model was considered the best-fitting model for pairs that did not fit an IM model significantly better. Conversely, pairs that fit an IM model better than an SI model, but did not show significant support for an IIM or SC model over the IM model, were assigned the IM model as best-fitting. Pairs that fit an IIM model significantly better than an IM model—and for which the IM model fit significantly better than the SI model—were assigned the IIM model as best-fitting. However, only one pair fit an IIM model better than an IM model.

To assign pairs to the secondary contact (SC) model—which is both more complex than the IM model and non-nested within it—we accepted it as the best-fitting model only when relative model support (ΔlnCL) exceeded the critical value for SI vs. SC (ΔlnCL > 5.991, 2 d.f.) **and** showed greater relative support than when compared against the IM model (ΔlnCL > 3.841, 1 d.f.). We confirmed that pairs best fitting the SC model displayed a higher frequency of blocks with few or no differences, consistent with a signal of recent gene flow (Supplementary Fig. 15).

### *1.5 Simulations – examining the effect of realistic recombination rates on model fit.*

To understand how our inference based on the S distribution is impacted by recombination within blocks and to test whether the pervasive signal of long term gene flow could be an artefact of ignoring recombination within blocks, we used *msprime* (*27*, *28*) to simulate blocks of the same size as our empirical dataset (Supplementary Table 2) under the best fitting SI history for each species pair. For every taxon pair we simulated data without recombination and with recombination assuming a fixed rate of $\rho$ = 1.03 cM/Mb, half the average recombination in *D. melanogaster* females to account for the achiasmy in males (*26*). Additionally, we assumed a per generation mutation rate of $3.32 \times 10^{-9}$ as mentioned above. For every pair, we simulated 2,000 blocks and normalised the S distributions as described above.

To quantify the bias in MLEs of parameters and to determine the false positive rate of model selection with and without recombination, we fitted simulated S distributions for every pair to the SI and IM models and assessed the difference in log-likelihood as detailed above for the empirical data. We simulated two pseudo-observed datasets (PODS): (1) without recombination and (2) with a realistic recombination rate and asked i) whether there is significant support for an IM relative to an SI model and ii) how rates of gene flow inferred from simulated data compare to empirical estimates.

When simulating blockwise data under an SI history and no recombination, parameter estimates closely match the true values and none of the PODS (0 out 93 species pairs) erroneously fit an IM model over a SI model (Supplementary Figure 8). This is consistent with the expected false positive rate (FPR) given the $\chi^2$ test we performed on ΔlnL. In contrast, when simulating under a SI model with recombination, 24 out of 93 PODS erroneously support an IM over an SI model which corresponds to a FPR of 25%. However, comparing the relative support (ΔlnL) between SI and IM for PODS (simulated under SI with recombination) with the empirical estimates shows that the IM model has considerably higher support in the real data compared to the PODS simulated without gene flow (two-sample Kolmogorov-Smirnov test: D = 0.79, p = <0.001).

Furthermore, we investigate the impact of recombination on our inference by comparing scaled rate of gene flow (M) and divergence time ($T_0$) estimates for PODS simulated under a SI model with recombination, with empirical parameter estimates. In the 77 species pairs that best fit an IM model in the empirical data the M estimates for the corresponding PODS (simulated with recombination under SI) are much lower than the empirical estimates (two-sample Kolmogorov-Smirnov test: D = 0.45, p= <0.001). Similarly, estimates for $T_0$ (two-sample Kolmogorov-Smirnov test: D = 0.58, p= <0.001) are also considerably lower for data simulated under the best fitting SI model compared to the empirical estimates across the 77 pairs. Thus, we conclude that while plausible rates of recombination may result in erroneous inference of gene flow, this effect is subtle and can neither explain the amount of gene flow we infer, nor the strength of support for speciation histories that involve gene flow that we find in the empirical data.

### 1.6 Understanding the effect of non-independent pairwise sampling on modelling gene flow.

To maximise the overlap with seminal comparative surveys of RI, our dataset includes non-independent species pairs. Assessing the impact of phylogenetically non-independent sampling on our main result is important because historic gene flow between a single pair of ancestral taxa may contribute to gene flow signals between in several focal species pairs. To understand the effect of the pseudoreplication, we used a pruned phylogeny of *Drosophila* inferred using single-copy orthologues from Kim et al. (2024) *(31)*, to trace each pairwise comparison of species back to its most recent common ancestor in the species tree. We quantified the number of pairwise comparisons that could be traced back to each speciation event (Supplementary Figure 11) and quantified average support for alternative models by averaging the difference in log-likelihood between nested models across all pairs corresponding to each node. Additionally, we also quantified node-level estimates for gene flow, divergence time and $N_e$ by averaging parameter estimates for all pairs assigned to an ancestral node. This simple node-level correction approach reduced our dataset to 36 observations across the *Drosophila* phylogeny.

Of the 36 node-level observations across our dataset, 31 nodes show evidence of significant support for an IM model over a strict isolation model (Supplementary Figure 12). Similarly, of the 36 nodes in the phylogenetically corrected dataset, 31 nodes have non-zero probabilities of migration (mean probability of migration = 0.68). Mean estimates of ancestral effective population ($N_e$ = 858,000) and divergence time ($T_0$ = 5,964,000) in our phylogenetically corrected dataset strongly resemble mean scaled effective population size ($N_e$ = 888,000) and divergence time ($T$ = 4,917,000) estimates. This suggests that the overwhelming signal of gene flow identified across pairwise comparisons is not inflated because of non-independent sampling. Additionally, in the corrected dataset, we found only one node (*D. athabasca – D. affinis*) where the IIM model fits significantly better than an IM model. However, we note that at some ancestral nodes, node-based estimates are heavily influenced by one or two species (e.g., *D. melanogaster* in the *melanogaster* group). Importantly, this not only reflects a bias in our genomic dataset but in the original comparative surveys of reproductive isolation which motivated our choice of taxa.

#### 1.7 Sensitivity analyses

For a number of pairs best fitting an IM model, parameter estimates for migration (M) and divergence time ($T_0$) hit the upper bounds, implying that for these pairs parameter estimates are unreliable. To assess the robustness of our main results, we removed pairs for which M was found to be larger than 10 (n=8) and $T_0$ larger than 9 (n=14), resulting in a reduced dataset of 71 pairs. We assessed again whether there was a difference in the probability of gene flow (two-sample Wilcoxon tests: W = 603, p = 0.847) and scaled divergence time (two-sample Wilcoxon tests: W = 683, p = 0.471) between sympatric and allopatric pairs with this reduced dataset only containing taxon pairs with reliable parameter estimates. As with our full dataset, we found no difference in the probability of migration and divergence times between allopatric and sympatric pairs.

**Supplementary Figures**

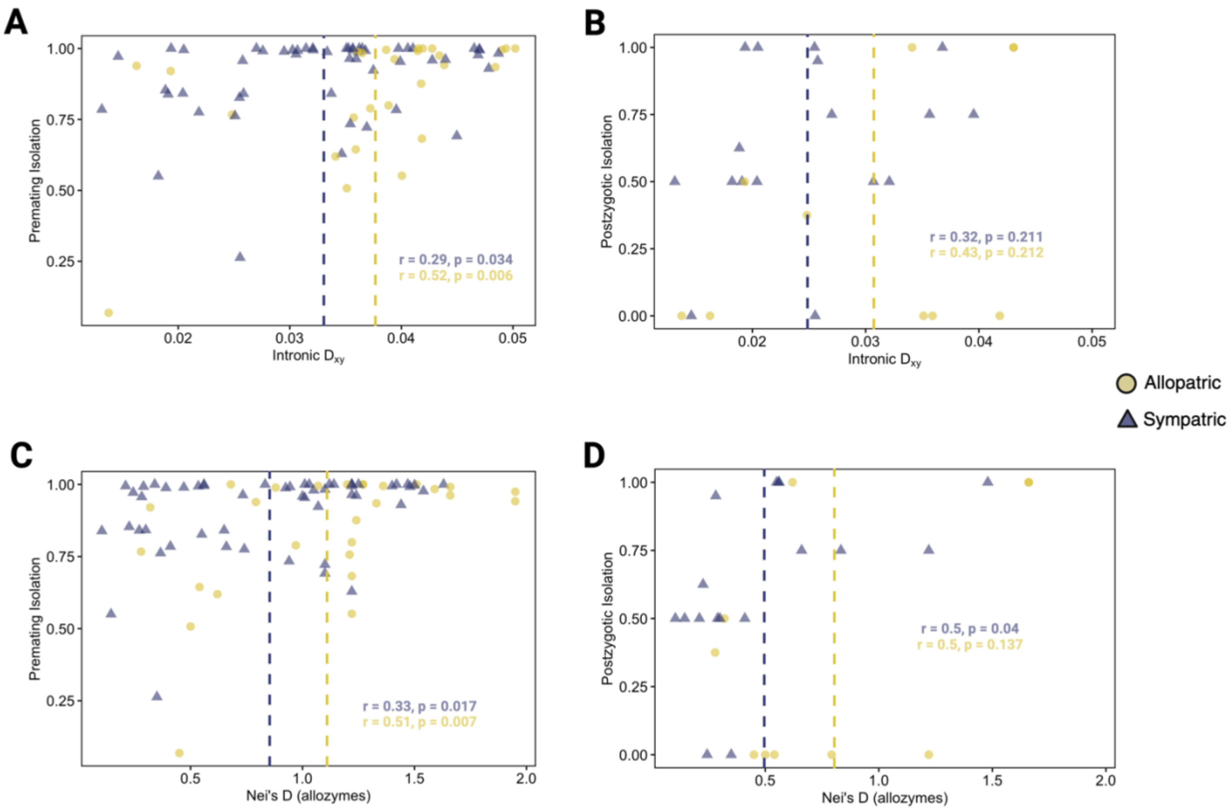


**Supplementary Figure 1:** Relationship between $d_{xy}$ (absolute genetic divergence in filtered introns) and measures of reproductive isolation. Genetic divergence calculated based on short intronic blocks sampled from whole-genome data in this study (A and B), and previous estimates of genetic divergence (Nei's D) based on allozymes (C and D). Dashed lines on the top two panels indicate mean intronic $d_{xy}$ (A and B) for allopatric and sympatric species pairs, and dashed lines in the bottom two panels indicate mean Nei's D calculated from allozymes for allopatric and sympatric pairs. A and C show estimates of premating isolation on the y-axis, and B and D show estimates of postzygotic isolation taken from previous comparative surveys of *Drosophila*. Pearson correlation coefficients and p-values for the relationship between measures of RI and measures of genetic divergence for sympatric and allopatric pairs are shown in each panel.

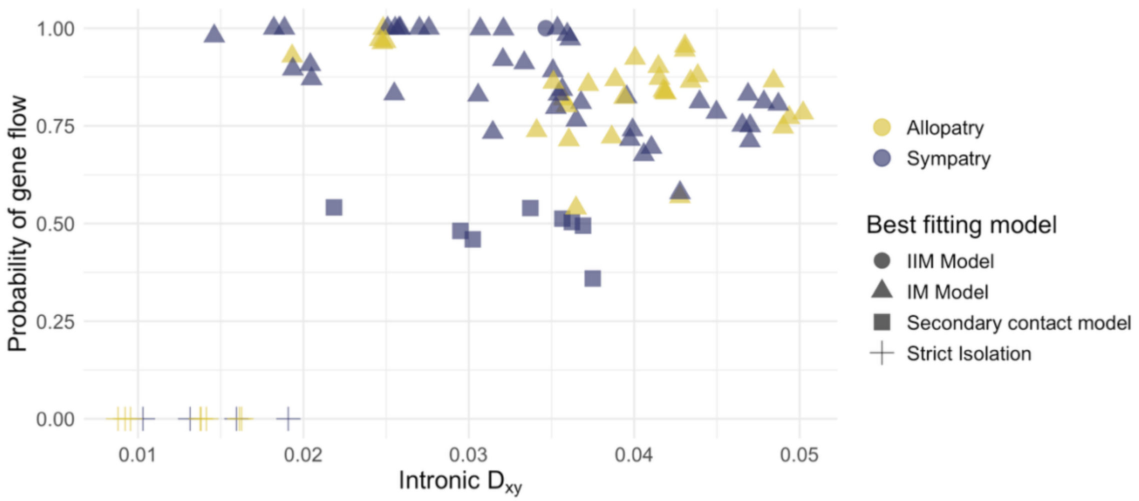


**Supplementary Figure 2:** Relationship between $d_{xy}$ (absolute genetic divergence in filtered introns) and probability of gene flow based on the best-fitting model. Pairs either fit an IM model with continuous gene flow, a secondary contact model with a burst of gene flow or a strict isolation model best (denoted by different shapes). Biogeography (whether pairs are currently allopatric or sympatric) is denoted by colours. The MIG model contains an infinite period of continuous gene flow and no definable divergence parameter, thus it is not possible calculate a probability of long-term migration for the one pair that showed best support for the MIG model.

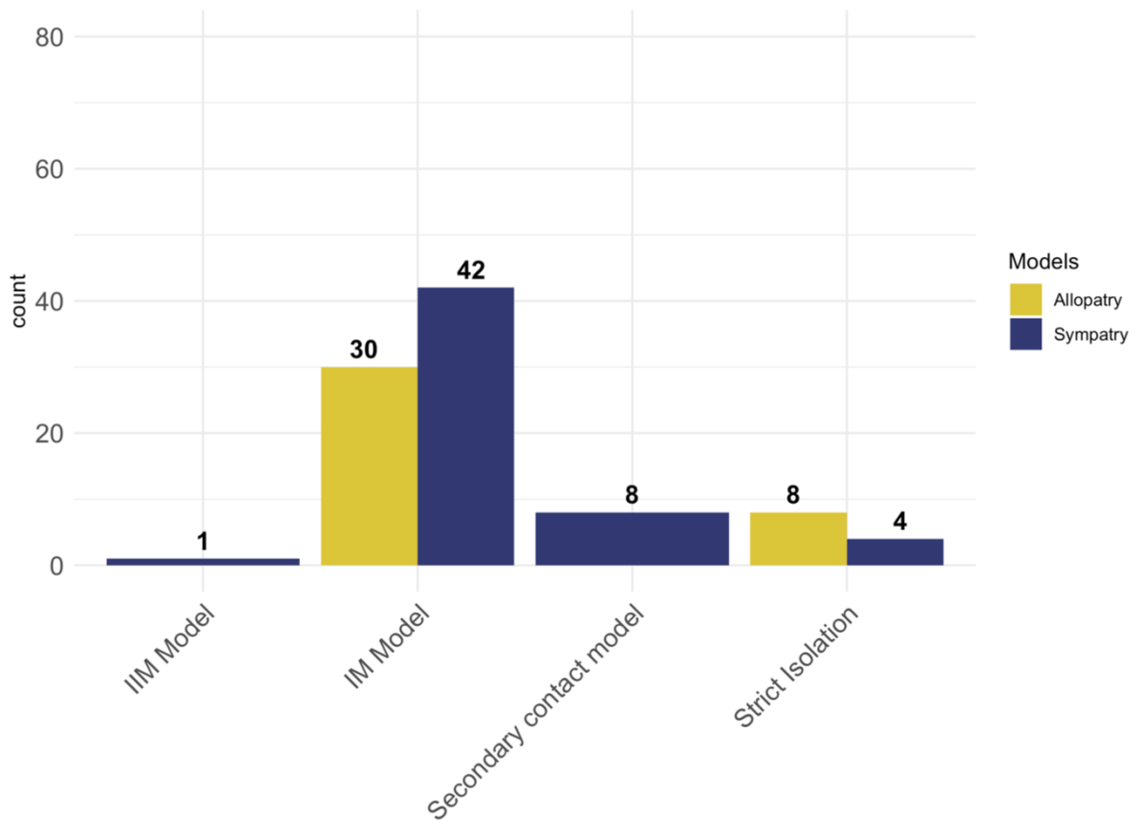


**Supplementary Figure 3:** Pairs and their best-fitting models. Pairs categorised into allopatry and sympatry, and the values on top of bars represent the number of allopatric and sympatric pairs best-fitting one of the three models. We found no allopatric pairs that best fit an SC or IIM model, so these are not represented here.

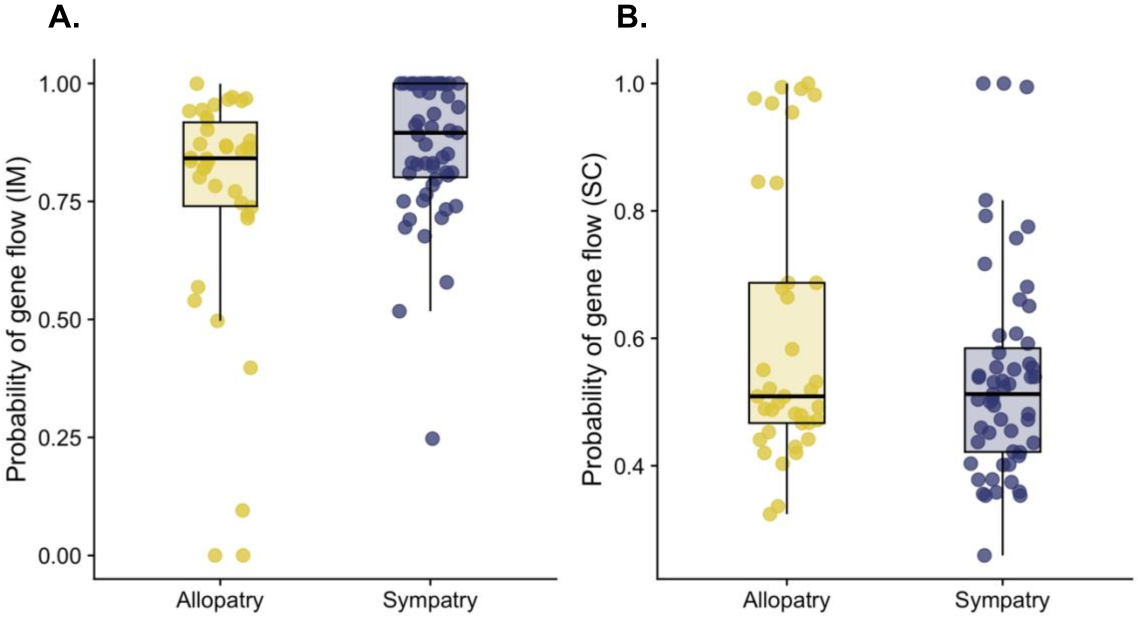


**Supplementary Figure 4:** Probability of gene flow under different demographic histories of species divergence for all pairs. For A and B, plots compare the probability of a single lineage migrating. A) Sympatric pairs are inferred to have a marginally higher mean probability of gene flow during a continuous migration phase than allopatric pairs under an IM model. B. Similar levels of inferred gene flow are seen during a short admixture phase under a secondary contact scenario.

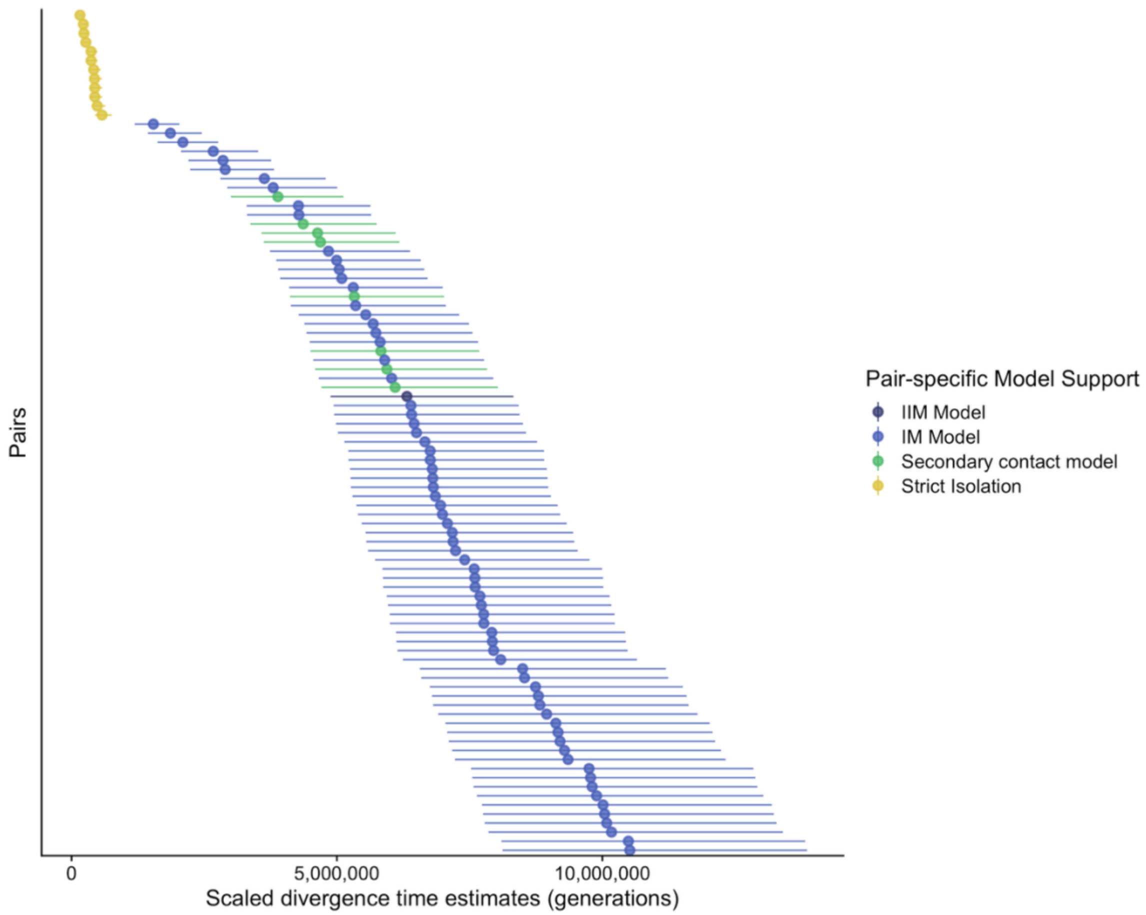


**Supplementary Figure 5:** Scaled initial divergence time estimates for *Drosophila* pairs using the best-fitting model. Parameter estimates were scaled (to number of generations) using a *D. melanogaster* mutation rate of 3.32 x $10^{-9}$ (*26*), represented by the circles. Upper and lower bound of divergence time for each pair reflect the upper and lower bounds of the *D. melanogaster* mutation rate (upper: 2.52 x $10^{-9}$; lower: 4.30 x $10^{-9}$). Pairs are coloured by their best-fitting model.

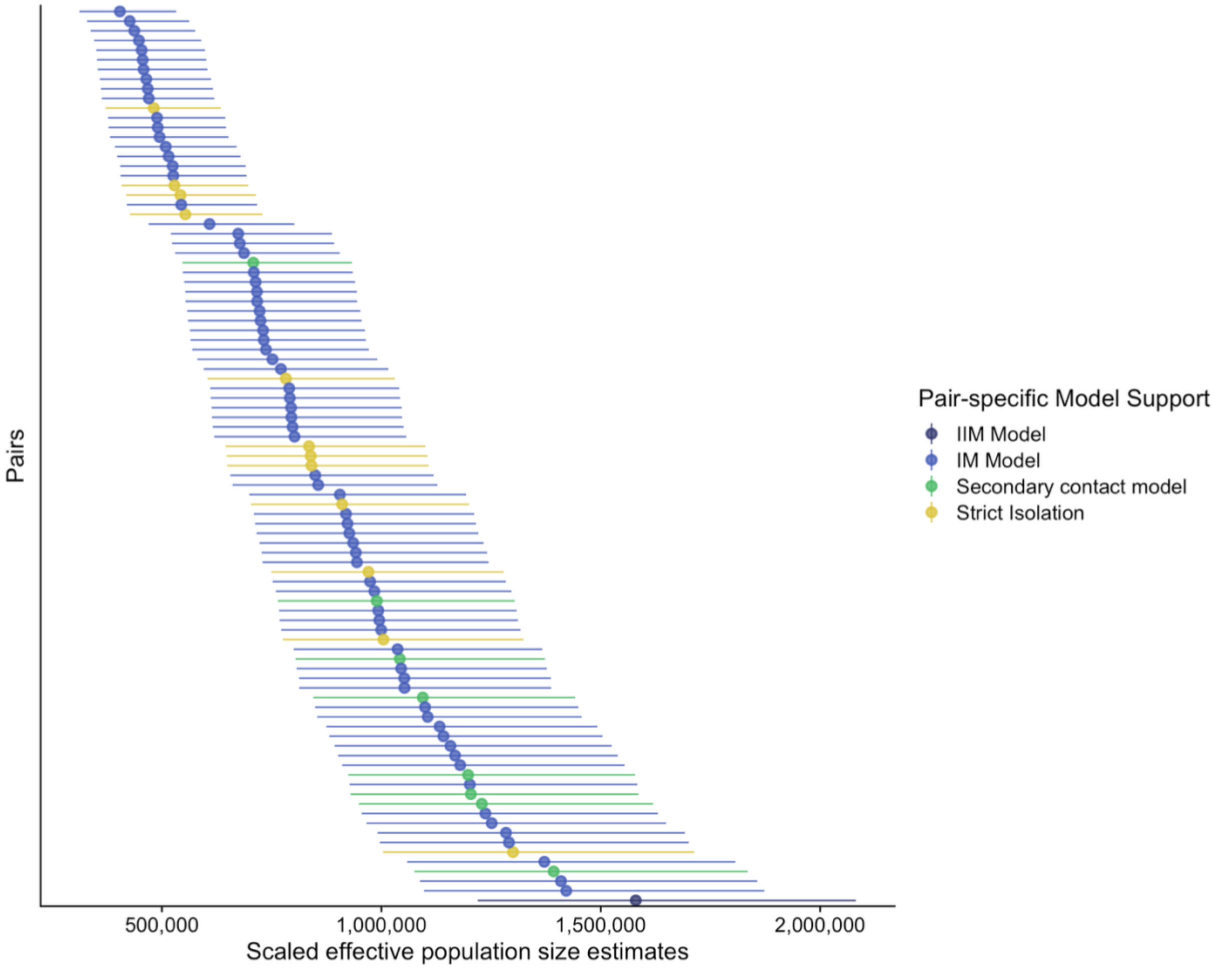


**Supplementary Figure 6:** Estimates of the effective population size ($N_e$) for Drosophila pairs under the best-fitting model. Parameter estimates were scaled using a *D. melanogaster* mutation rate of 3.32 x $10^{-9}$ (*26*). Upper and lower bounds reflect the upper and lower bounds of the *D. melanogaster* mutation rate (upper: 2.52 x $10^{-9}$; lower: 4.30 x $10^{-9}$). Pairs are coloured by their best-fitting model.

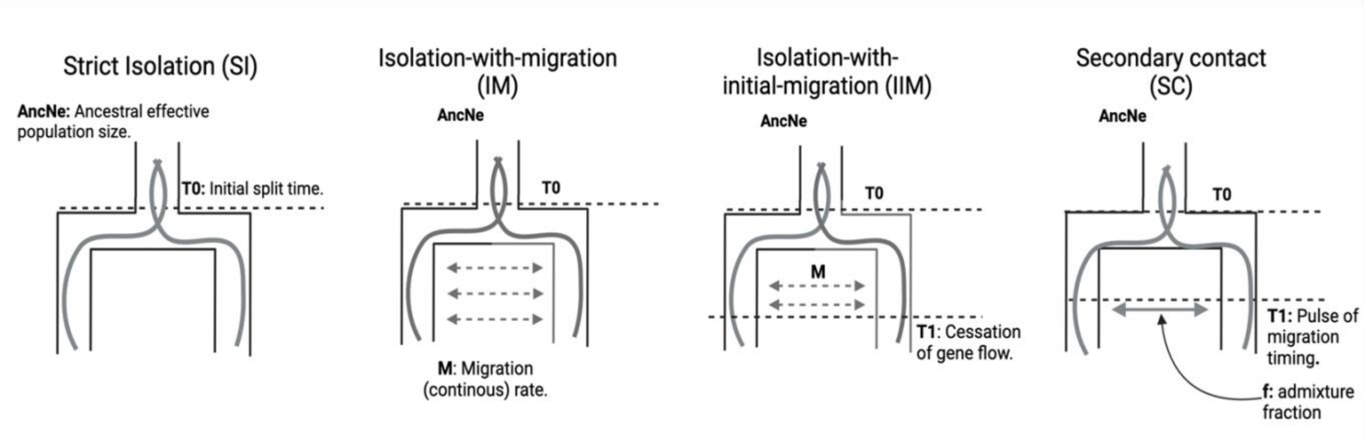


**Supplementary Figure 7:** Speciation models considered in the analysis. Models include a Strict Isolation (SI) model representing allopatric speciation without any gene flow (2 parameters), an isolation-with-migration (IM) model with continuous gene flow (3 parameters), an isolation-with-initial-migration model (IIM) which includes an initial phase of migration followed by no gene flow (4 parameters), and a secondary contact (SC) model which includes a recent pulse of migration (4 parameters).

276

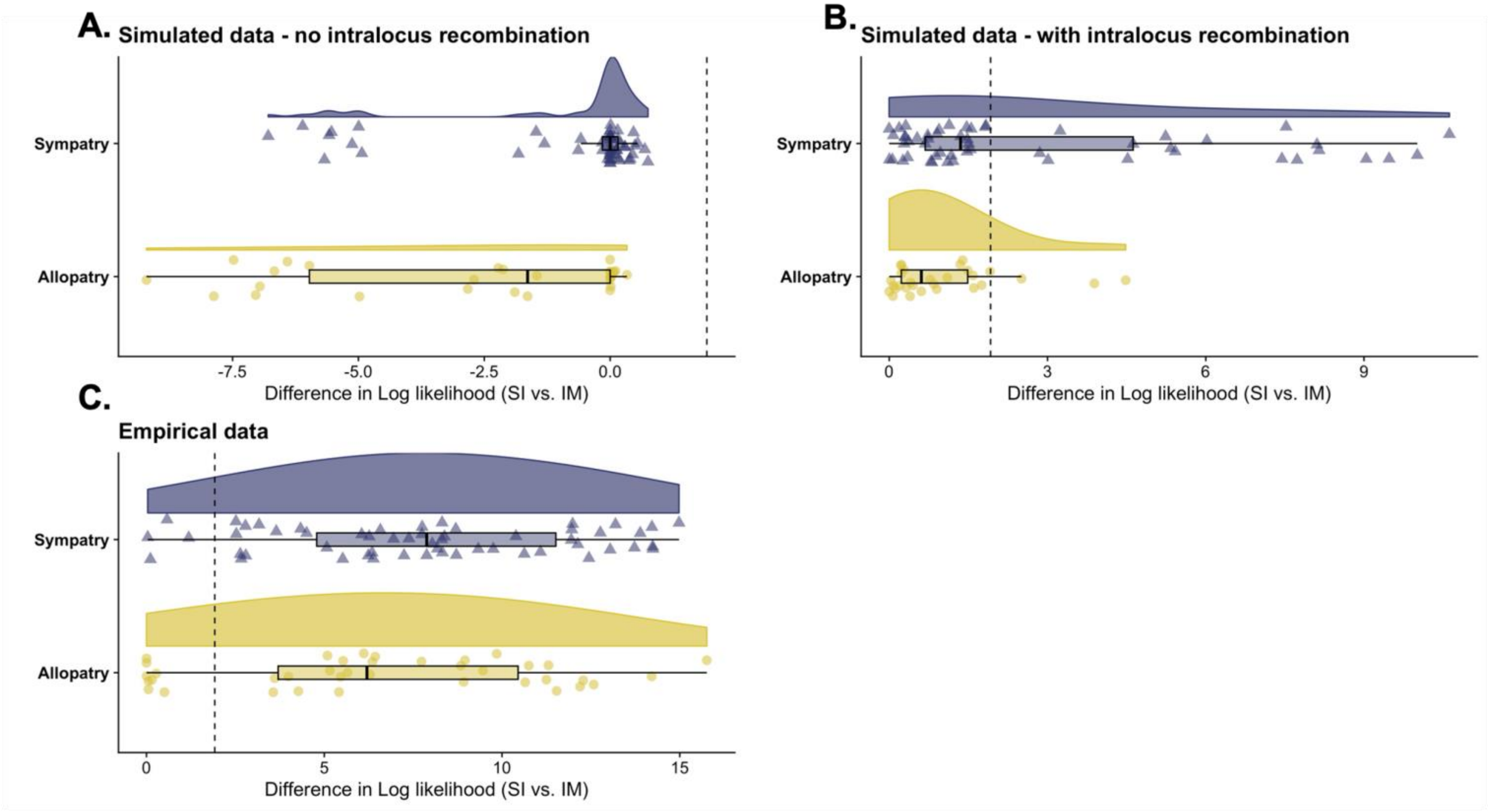


**Supplementary Figure 8:** Comparison of the difference in log-likelihood between a strict isolation (SI) model and an isolation-with-migration model (IM) between **A.** Data simulated according to optimized strict divergence history parameter estimates but with no recombination, **B.** Data simulated under an optimized strict divergence history with D. melanogaster levels of recombination, and **C.** empirical data. The dashed line indicates the critical threshold ($p < 0.05$). **A.** Simulations with no recombination show a false positive rate of detecting a history of speciation-with-gene-flow where there is none is 0%. **B.** Simulated data with realistic levels of recombination show a higher false positive rate of 25%. Comparison of **B.** and **C.** demonstrate that simulations with recombination do not recapitulate as strong support for a speciation history with gene flow across allopatric and sympatric pairs as the empirical data.

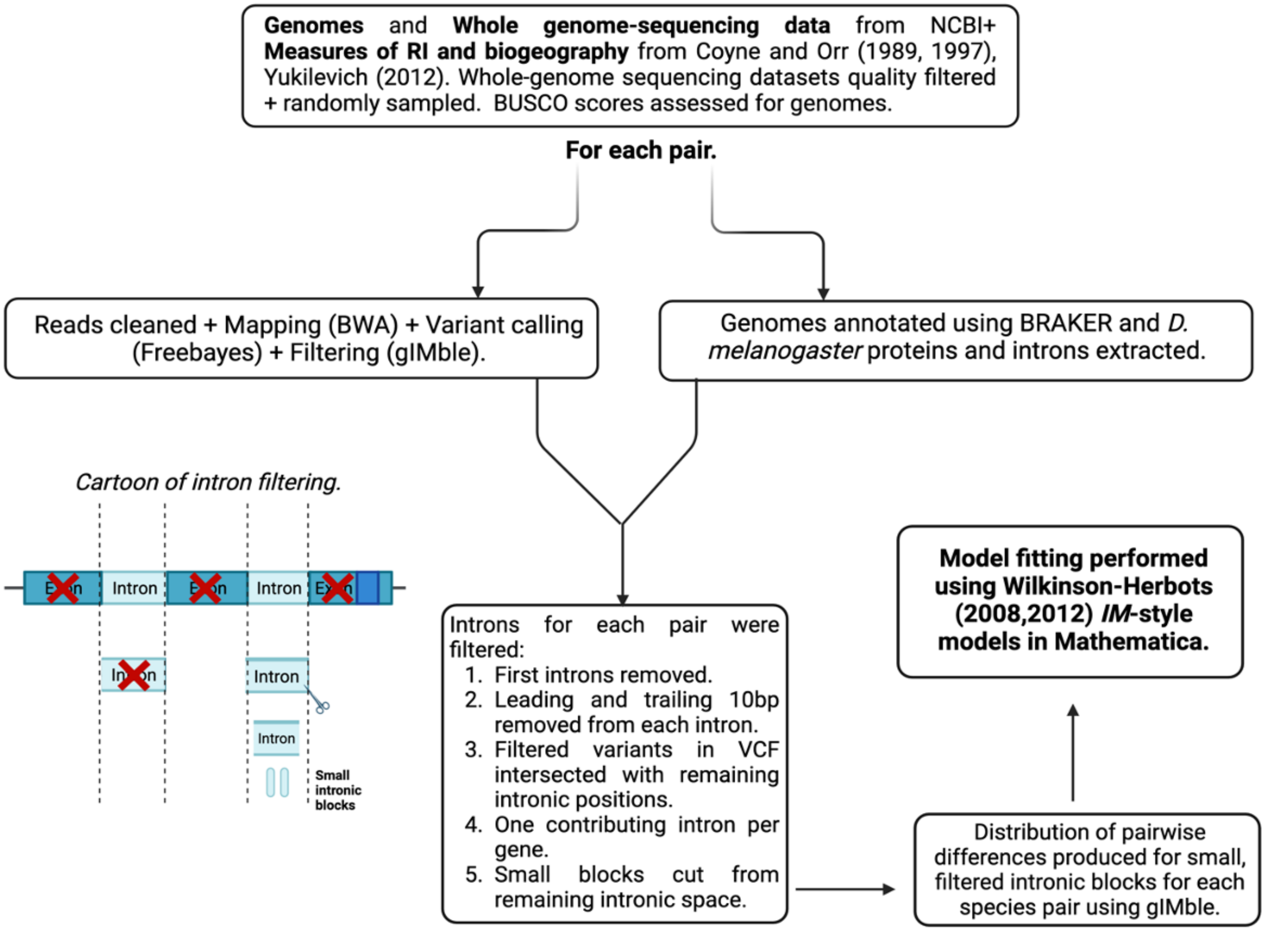


**Supplementary Figure 9:** Flowchart of analysis pipeline. Describing workflow for the analysis beginning with gathering data on reproductive isolation and biogeography for Drosophila pairs from previous comparative analyses, and whole-genome sequencing datasets and genomes from NCBI, and ending with model fitting for remaining pairs. Intron filtering cartoon alongside the description highlights main filtering decisions taken to produce the datasets used for the model fitting.

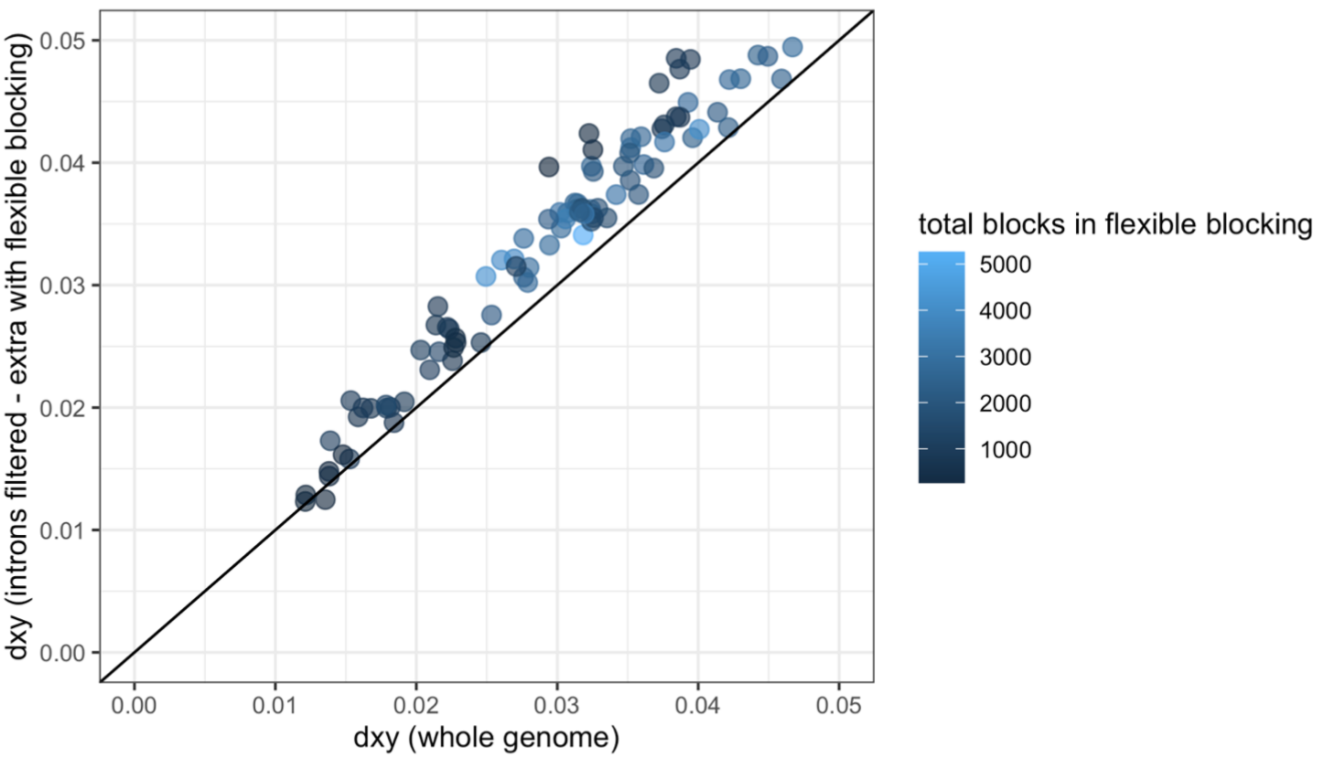


**Supplementary Figure 10:** The relationship between $d_{XY}$ calculated in fixed 200bp blocks across the whole genome and $d_{XY}$ calculated in filtered, intronic blocks across 93 species pairs. Points are coloured based on how many intronic blocks remained and were used in the model fitting analysis for each pair.

.

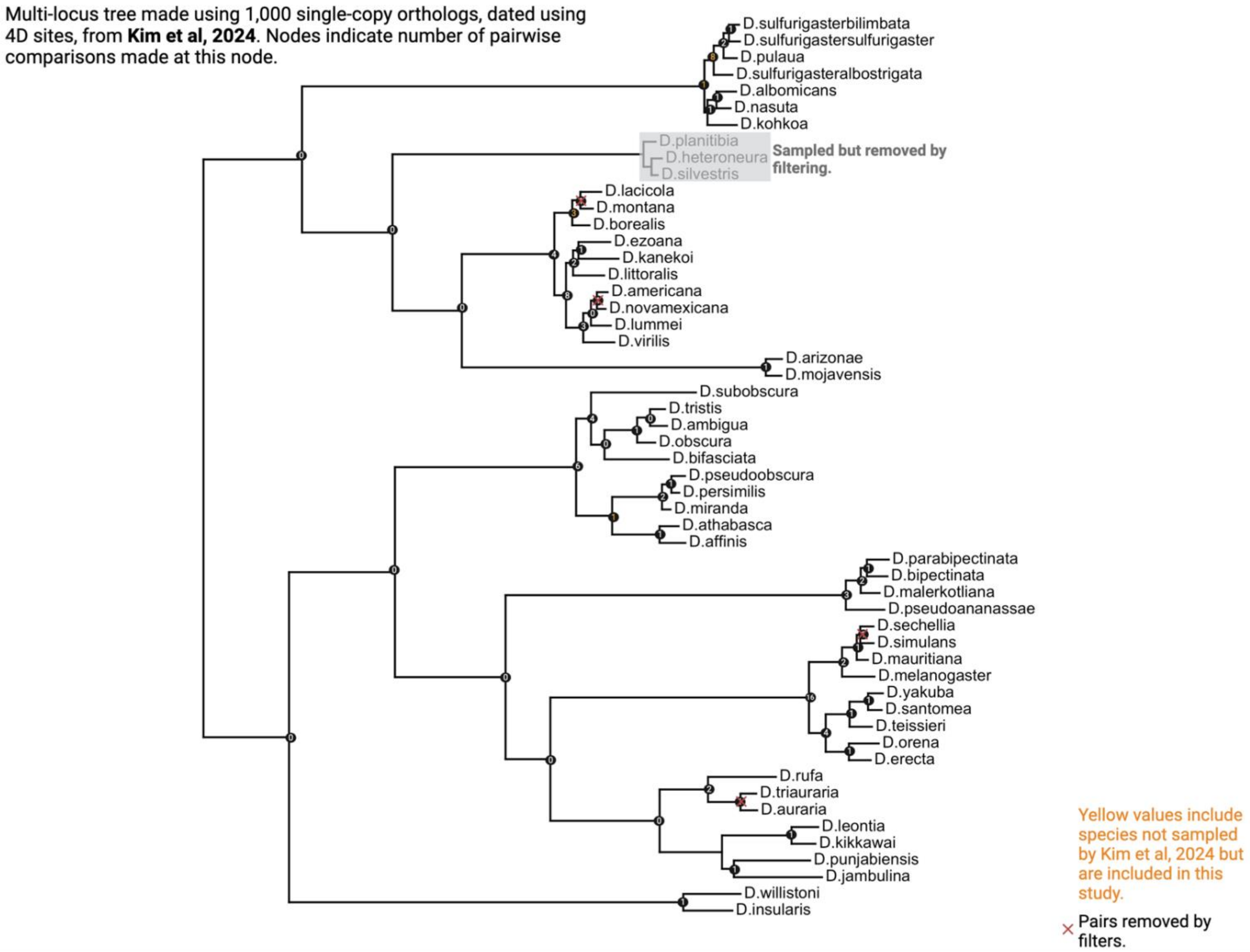


**Supplementary Figure 11:** Pruned phylogenetic tree of Drosophila taken from Kim et al (2024). Phylogeny contains species considered in this study. Numbers at each node represent the number of pairs that have diverged at these nodes. This is a visualisation representation of the level of non-independence in our dataset.

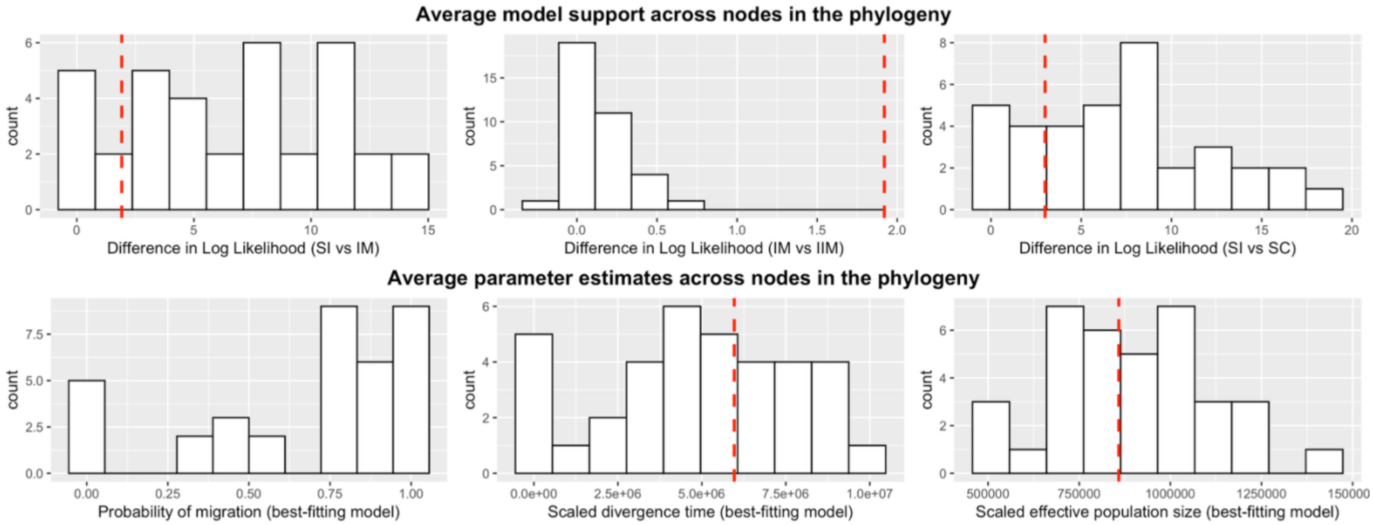


**Supplementary Figure 12:** Average model support and parameter estimates across nodes in the Drosophila phylogeny. Top panel shows averaged difference in log likelihood across nodes in the phylogeny for nested model comparisons. The red dashed line indicates the critical threshold ($p < 0.05$). The black dashed line denotes 0, where positive values indicate nodes with average support for an IIM model over the SC model, and vice versa. Bottom panel shows average optimised parameter estimates according to the best-fitting model in pairwise comparisons, across nodes in the phylogeny.

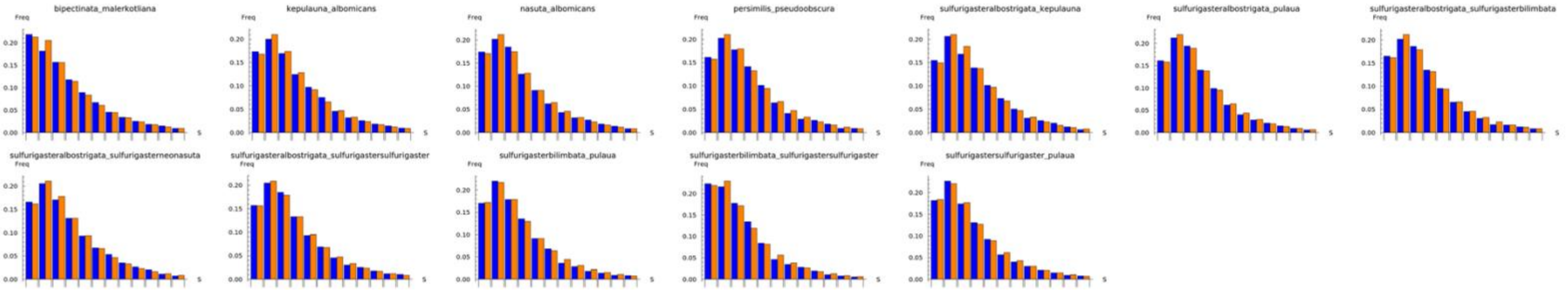


**Supplementary Figure 13:** Observed (blue) versus expected (orange) S distributions for pairs best fitting a Strict Isolation (SI) model.

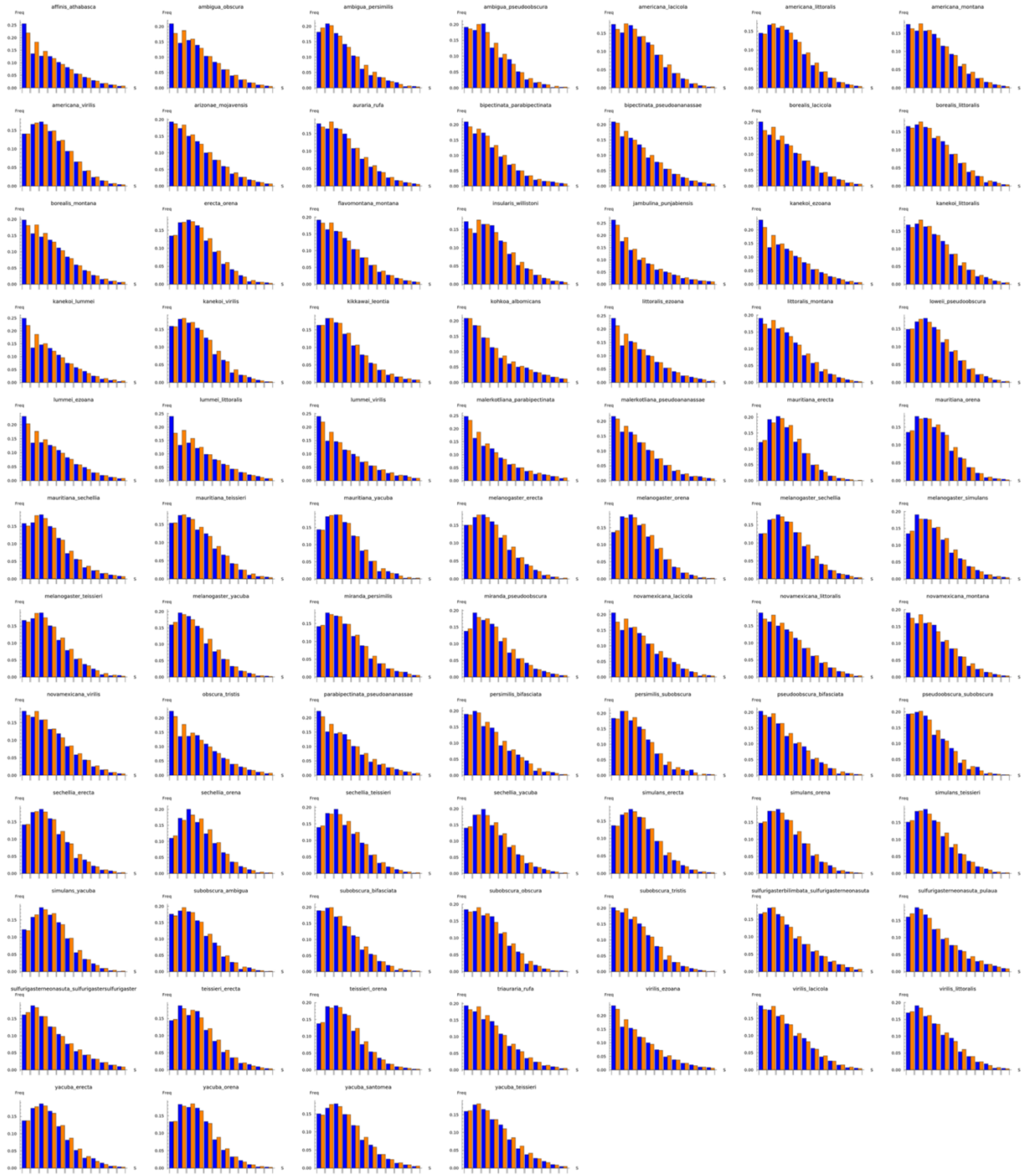


**Supplementary Figure 14:** Observed (blue) versus expected (orange) S distributions for pairs that fit an IM model significantly better than SI model. Pairs which include red stars best-fit an SC model, and only one pair, with a green star, best-fits the IIM model.

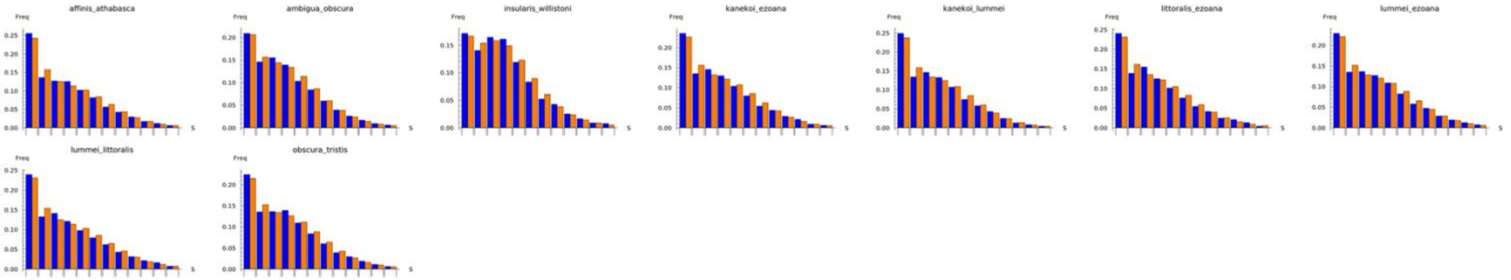


**Supplementary Figure 15:** Observed (blue) versus expected (orange) S distributions for pairs best fitting a Secondary contact (SC) model

***3.2 Tables: 1 - 6***

| Pair | Accession Species A | Accession Species B |
|---|---|---|
| *D. affinis - D. athabasca* | ERR127384 | SRR9967640 |
| *D. ambigua - D. obscura* | SRR13070667 | SRR13070671 |
| *D. ambigua - D. persimilis* | SRR12899691 | SRR13070667 |
| *D. ambigua - D. pseudoobscura* | SRR11230572 | SRR13070667 |
| *D. ambigua - D. tristis* | SRR13070667 | SRR13070669 |
| *D. americana - D. lacicola* | ERR2610695 | SRR20226091 |
| *D. americana - D. littoralis* | ERR2610695 | SRR13070670 |
| *D. americana - D. montana* | ERR2610695 | SRR3218635 |
| *D. americana - D. novamexicana* | ERR2610692 | ERR2610695 |
| *D. americana - D. virilis* | ERR2610695 | SRR9426114 |
| *D. arizonae - D. mojavensis* | SRR2070760 | SRR6425997 |
| *D. auraria - D. rufa* | SRR13070713 | SRR13188937 |
| *D. auraria - D. triauraria* | SRR13188922 | SRR13188937 |
| *D. bipectinata - D. malerkotliana (pallens)* | SRR13070716 | SRR15048573 |
| *D. bipectinata - D. parabipectinata* | SRR13070721 | SRR15048573 |
| *D. bipectinata - D. pseudoananassae (nigrens)* | SRR13070715 | SRR15048573 |
| *D. borealis - D. lacicola* | SRR20226101 | SRR20226091 |
| *D. borealis - D. littoralis* | SRR13070670 | SRR20226101 |
| *D. borealis - D. montana* | SRR3218635 | SRR20226101 |
| *D. erecta - D. orena* | SRR1977592 | SRR6425990 |
| *D. flavomontana - D. montana* | SRR3218635 | SRR20226092 |
| *D. heteroneura - D. planitibia* | SRR2233094 | SRR2233332 |
| *D. heteroneura - D. silvestris* | SRR2233094 | SRR2233314 |
| *D. insularis - D. willistoni* | SRR13070692 | SRR6426003 |
| *D. jambulina - D. punjabiensis* | SRR15048354 | SRR9997923 |

| | | |
|---|---|---|
| *D. kanekoi - D. ezoana* | SRR20226095 | SRR20226093 |
| *D. kanekoi - D. littoralis* | SRR13070670 | SRR20226093 |
| *D. kanekoi - D. virilis* | SRR9426114 | SRR20226093 |
| *D. kepulauna - D. albomicans* | DRR076000 | SRR10100373 |
| *D. kikkawai - D. leontia* | SRR15048447 | SRR1810494 |
| *D. kohkoa - D. albomicans* | DRR076000 | DRR160734 |
| *D. lacicola - D. montana* | SRR3218635 | SRR20226091 |
| *D. littoralis - D. ezoana* | SRR13070670 | SRR20226095 |
| *D. littoralis - D. montana* | SRR13070670 | SRR3218635 |
| *D. loweii - D. pseudoobscura* | SRR11230572 | SRR9967663 |
| *D. lummei - D. ezoana* | SRR5278982 | SRR20226095 |
| *D. lummei - D. littoralis* | SRR13070670 | SRR5278982 |
| *D. lummei - D. virilis* | SRR5278982 | SRR9426114 |
| *D. malerkotliana (pallens) - D. parabipectinata* | SRR13070716 | SRR13070721 |
| *D. malerkotliana (pallens) - D. pseudoananassae (nigrens)* | SRR13070715 | SRR13070716 |
| *D. mauritiana - D. erecta* | SRR1560109 | SRR6425990 |
| *D. mauritiana - D. orena* | SRR1560109 | SRR1977592 |
| *D. mauritiana - D. sechellia* | SRR14138507 | SRR1560109 |
| *D. mauritiana - D. teissieri* | SRR1560109 | SRR5860615 |
| *D. mauritiana - D. yakuba* | SRR1560109 | SRR9700025 |
| *D. melanogaster - D. erecta* | SRR3931586 | SRR6425990 |
| *D. melanogaster - D. orena* | SRR1977592 | SRR3931586 |
| *D. melanogaster - D. sechellia* | SRR14138507 | SRR3931586 |
| *D. melanogaster - D. simulans* | SRR11456787 | SRR3931586 |
| *D. melanogaster - D. teissieri* | SRR3931586 | SRR5860615 |
| *D. melanogaster - D. yakuba* | SRR3931586 | SRR9700025 |

| | | |
|---|---|---|
| *D. miranda - D. bifasciata* | DRR061004 | SRR789669 |
| *D. miranda - D. persimilis* | SRR12899691 | SRR789669 |
| *D. miranda - D. pseudoobscura* | SRR11230572 | SRR789669 |
| *D. miranda - D. subobscura* | SRR789669 | SRR9967664 |
| *D. nasuta - D. albomicans* | DRR076000 | SRR10100355 |
| *D. novamexicana - D. lacicola* | ERR2610692 | SRR20226091 |
| *D. novamexicana - D. littoralis* | ERR2610692 | SRR13070670 |
| *D. novamexicana - D. montana* | ERR2610692 | SRR3218635 |
| *D. novamexicana - D. virilis* | ERR2610692 | SRR9426114 |
| *D. obscura - D. tristis* | SRR13070669 | SRR13070671 |
| *D. parabipectinata - D. pseudoananassae* | SRR13070715 | SRR13070721 |
| *D. persimilis - D. bifasciata* | DRR061004 | SRR12899691 |
| *D. persimilis - D. pseudoobscura* | SRR11230572 | SRR12899691 |
| *D. persimilis - D. subobscura* | SRR12899691 | SRR9967664 |
| *D. planitibia - D. silvestris* | SRR2233314 | SRR2233332 |
| *D. pseudoobscura - D. bifasciata* | DRR061004 | SRR11230572 |
| *D. pseudoobscura - D. subobscura* | SRR11230572 | SRR9967664 |
| *D. sechellia - D. erecta* | SRR14138507 | SRR6425990 |
| *D. sechellia - D. orena* | SRR14138507 | SRR1977592 |
| *D. sechellia - D. teissieri* | SRR14138507 | SRR5860615 |
| *D. sechellia - D. yakuba* | SRR14138507 | SRR9700025 |
| *D. simulans - D. erecta* | SRR11456787 | SRR6425990 |
| *D. simulans - D. orena* | SRR11456787 | SRR1977592 |
| *D. simulans - D. sechellia* | SRR11456787 | SRR14138507 |
| *D. simulans -D. teissieri* | SRR11456787 | SRR5860615 |
| *D. simulans - D. yakuba* | SRR11456787 | SRR9700025 |
| *D. subobscura - D. ambigua* | SRR13070667 | SRR9967664 |

| | | |
|---|---|---|
| *D. subobscura - D. bifasciata* | DRR061004 | SRR9967664 |
| *D. subobscura - D. obscura* | SRR13070671 | SRR9967664 |
| *D. subobscura - D. tristis* | SRR13070669 | SRR9967664 |
| *D. sulfurigaster albostrigata - D. kepulauna* | SRR10100373 | SRR10100407 |
| *D. sulfurigaster albostrigata - D. pulaua* | SRR10100335 | SRR10100407 |
| *D. sulfurigaster albostrigata - D. sulfurigaster bilimbata* | SRR10100308 | SRR10100323 |
| *D. sulfurigaster albostrigata - D. sulfurigaster neonasuta* | SRR10100407 | SRR5059342 |
| *D. sulfurigaster albostrigata - D. sulfurigaster sulfurigaster* | SRR10100323 | SRR10100401 |
| *D. sulfurigaster bilimbata - pulaua* | SRR10100308 | SRR10100335 |
| *D. sulfurigaster bilimbata - sulfurigaster neonasuta* | SRR10100308 | SRR5059342 |
| *D. sulfurigaster bilimbata - D. sulfurigaster sulfurigaster* | SRR10100308 | SRR10100401 |
| *D. sulfurigaster neonasuta - pulaua* | SRR10100335 | SRR5059342 |
| *D. sulfurigaster neonasuta - D. sulfurigaster sulfurigaster* | SRR10100401 | SRR5059342 |
| *D. sulfurigaster sulfurigaster - D. pulaua* | SRR10100335 | SRR10100401 |
| *D. teissieri - D. erecta* | SRR5860615 | SRR6425990 |
| *D. teissieri - D. orena* | SRR1977592 | SRR5860615 |
| *D. triauraria - D. rufa* | SRR13070713 | SRR13188922 |
| *D. virilis - D. ezoana* | SRR9426114 | SRR20226095 |
| *D. virilis - D. lacicola* | SRR9426114 | SRR20226091 |
| *D. virilis - D. littoralis* | SRR13070670 | SRR9426114 |
| *D. yakuba - D. erecta* | SRR6425990 | SRR9700025 |
| *D. yakuba - D. orena* | SRR1977592 | SRR9700025 |
| *D. yakuba - D. santomea* | SRR5860642 | SRR9700025 |

| | | |
|---|---|---|
| *D. yakuba - D. teissieri* | SRR5860615 | SRR9700025 |

**Supplementary Table 1:** Species pairs and SRA accessions for whole-genome sequencing samples used in this study.

| Pair | Intronic $D_{XY}$ | No. of blocks | Block size |
| --- | --- | --- | --- |
| *D. affinis - D. athabasca* | 0.028 | 1300 | 137 |
| *D. ambigua - D. obscura* | 0.0119 | 2460 | 99 |
| *D. ambigua - D. persimilis* | 0.0128 | 738 | 70 |
| *D. ambigua - D. pseudoobscura* | 0.0126 | 725 | 70 |
| *D. americana - D. lacicola* | 0.0128 | 2130 | 82 |
| *D. americana - D. littoralis* | 0.0132 | 2230 | 84 |
| *D. americana - D. montana* | 0.0115 | 2140 | 82 |
| *D. americana - D. virilis* | 0.0129 | 2530 | 84 |
| *D. arizonae - D. mojavensis* | 0.0145 | 3120 | 121 |
| *D. auraria- D. rufa* | 0.0132 | 2080 | 83 |
| *D. bipectinata - D. malerkotliana* | 0.0108 | 2670 | 157 |
| *D. bipectinata - D. parabipectinata* | 0.0116 | 2660 | 165 |
| *D. bipectinata - D. pseudoananassae* | 0.0117 | 1920 | 116 |
| *D. borealis - D. lacicola* | 0.0133 | 3270 | 94 |
| *D. borealis - D. littoralis* | 0.0139 | 2070 | 72 |
| *D. borealis - D. montana* | 0.01 | 3280 | 93 |
| *D. erecta - D. orena* | 0.0143 | 2440 | 95 |
| *D. flavomontana- D. montana* | 0.0103 | 3460 | 98 |
| *D. insularis - D. willistoni* | 0.0147 | 1720 | 80 |
| *D. jambulina - D. punjabiensis* | 0.0122 | 2590 | 109 |
| *D. kanekoi - D. ezoana* | 0.0138 | 2450 | 84 |
| *D. kanekoi - D. littoralis* | 0.0144 | 2160 | 75 |
| *D. kanekoi - D. lummei* | 0.012 | 2230 | 81 |
| *D. kanekoi - D. virilis* | 0.0141 | 1890 | 67 |

| | | | |
|---|---|---|---|
| *D. kepulauna - D. albomicans* | 0.0125 | 3510 | 185 |
| *D. kikkawai- D. leontia* | 0.00865 | 3010 | 205 |
| *D. kohkoa - D. albomicans* | 0.0128 | 3790 | 111 |
| *D. littoralis - D. ezoana* | 0.0131 | 2480 | 89 |
| *D. littoralis - D. montana* | 0.0118 | 2160 | 76 |
| *D. loweii - D. pseudoobscura* | 0.0135 | 2150 | 98 |
| *D. lummei - D. ezoana* | 0.0125 | 2450 | 83 |
| *D. lummei - D. littoralis* | 0.0125 | 2350 | 87 |
| *D. lummei - D. virilis* | 0.0107 | 2680 | 117 |
| *D. malerkotliana - D. parabipectinata* | 0.0139 | 2950 | 159 |
| *D. malerkotliana - D. pseudoananassae* | 0.0123 | 1950 | 116 |
| *D. mauritiana - D. erecta* | 0.0138 | 1580 | 82 |
| *D. mauritiana - D. orena* | 0.0143 | 1590 | 78 |
| *D. mauritiana - D. sechellia* | 0.0148 | 3140 | 155 |
| *D. mauritiana - D. teissieri* | 0.0139 | 1930 | 72 |
| *D. mauritiana - D. yakuba* | 0.0132 | 1400 | 83 |
| *D. melanogaster - D. erecta* | 0.0141 | 1430 | 85 |
| *D. melanogaster - D. orena* | 0.0135 | 3990 | 73 |
| *D. melanogaster - D. sechellia* | 0.0148 | 6390 | 88 |
| *D. melanogaster - D. simulans* | 0.011 | 2010 | 118 |
| *D. melanogaster - D. teissieri* | 0.0123 | 1300 | 86 |
| *D. melanogaster - D. yakuba* | 0.0126 | 1180 | 85 |
| *D. miranda - D. persimilis* | 0.0149 | 2670 | 155 |
| *D. miranda - D. pseudoobscura* | 0.0146 | 2560 | 146 |
| *D. nasuta - D. albomicans* | 0.0143 | 3350 | 218 |
| *D. novamexicana - D. lacicola* | 0.0142 | 2430 | 72 |

| | | | |
|---|---|---|---|
| *D. novamexicana - D. littoralis* | 0.0144 | 2560 | 81 |
| *D. novamexicana - D. montana* | 0.0128 | 2620 | 84 |
| *D. novamexicana - D. virilis* | 0.0146 | 2810 | 85 |
| *D. obscura - D. tristis* | 0.0123 | 2530 | 102 |
| *D. parabipectinata - D. pseudoananassae* | 0.0139 | 2070 | 120 |
| *D. persimilis - D. bifasciata* | 0.0133 | 682 | 69 |
| *D. persimilis - D. pseudoobscura* | 0.0143 | 2700 | 228 |
| *D. persimilis - D. subobscura* | 0.0124 | 377 | 72 |
| *D. pseudoobscura - D. bifasciata* | 0.0131 | 703 | 68 |
| *D. pseudoobscura - D. subobscura* | 0.0123 | 367 | 76 |
| *D. sechellia - D. erecta* | 0.0144 | 1800 | 60 |
| *D. sechellia - D. orena* | 0.0147 | 4900 | 70 |
| *D. sechellia - D. teissieri* | 0.0133 | 1660 | 61 |
| *D. sechellia - D. yakuba* | 0.0139 | 1600 | 61 |
| *D. simulans - D. erecta* | 0.0134 | 1680 | 64 |
| *D. simulans - D. orena* | 0.0123 | 1470 | 75 |
| *D. simulans - D. teissieri* | 0.0119 | 1710 | 68 |
| *D. simulans - D. yakuba* | 0.013 | 1730 | 70 |
| *D. subobscura - D. ambigua* | 0.0122 | 673 | 63 |
| *D. subobscura - D. bifasciata* | 0.0125 | 669 | 62 |
| *D. subobscura - D. obscura* | 0.0124 | 658 | 64 |
| *D. subobscura - D. tristis* | 0.0118 | 643 | 62 |
| *D. sulfurigaster albostrigata - D. kepulauna* | 0.0106 | 2670 | 188 |
| *D. sulfurigaster albostrigata - D. pulaua* | 0.0102 | 1460 | 291 |
| *D. sulfurigaster albostrigata - D. sulfurigaster bilimbata* | 0.0125 | 2980 | 212 |

| | | | |
|---|---|---|---|
| *D. sulfurigaster albostrigata - D. sulfurigaster neonasuta* | 0.0111 | 2920 | 186 |
| *D. sulfurigaster albostrigata - D. sulfurigaster sulfurigaster* | 0.0126 | 2950 | 217 |
| *D. sulfurigaster bilimbata - D. pulaua* | 0.0131 | 1750 | 314 |
| *D. sulfurigaster bilimbata - D. sulfurigaster neonasuta* | 0.0137 | 3300 | 120 |
| *D. sulfurigaster bilimbata - D. sulfurigaster sulfurigaster* | 0.0117 | 1710 | 341 |
| *D. sulfurigaster neonasuta - D. pulaua* | 0.0137 | 3340 | 121 |
| *D. sulfurigaster neonasuta - D. sulfurigaster sulfurigaster* | 0.0138 | 3400 | 122 |
| *D. sulfurigaster sulfurigaster - D. pulaua* | 0.0125 | 1620 | 325 |
| *D. teissieri- D. erecta* | 0.0134 | 1850 | 64 |
| *D. teissieri - D. orena* | 0.0125 | 1640 | 75 |
| *D. triauraria - D. rufa* | 0.0126 | 1830 | 83 |
| *D. virilis - D. ezoana* | 0.0133 | 1930 | 85 |
| *D. virilis - D. lacicola* | 0.0142 | 1800 | 77 |
| *D. virilis - D. littoralis* | 0.014 | 1920 | 72 |
| *D. yakuba - D. erecta* | 0.014 | 1800 | 64 |
| *D. yakuba - D. orena* | 0.0136 | 1590 | 74 |
| *D. yakuba - D. santomea* | 0.0118 | 3050 | 147 |
| *D. yakuba - D. teissieri* | 0.0125 | 2440 | 90 |

**Supplementary Table 2:** Species pairs and summary statistics calculated from filtered intronic data. Includes estimates of intronic absolute genetic divergence ($D_{XY}$) for each sample, and the number and size of filtered intronic blocks used in the analysis.

| Pair | *Species (Genome)* | Source | Reference |
|---|---|---|---|
| D. affinis - D. athabasca | *D. athabasca* | SAMN12214156 | Bracewell, R., Chatla, K., Nalley, M.J. and Bachtrog, D., 2019. Dynamic turnover of centromeres drives karyotype evolution in Drosophila. *Elife*, *8*, p.e49002. |
| D. ambigua - D. obscura | *D. obscura* | SAMN16729671 | Kim, B.Y., Wang, J.R., Miller, D.E., Barmina, O., Delaney, E., Thompson, A., Comeault, A.A., Peede, D., D'Agostino, E.R., Pelaez, J. and Aguilar, J.M., 2021. Highly contiguous assemblies of 101 drosophilid genomes. *Elife*, *10*, p.e66405. |
| D. ambigua - D. persimilis | *D. persimilis* | SAMN16729701 | Kim, B.Y., Wang, J.R., Miller, D.E., Barmina, O., Delaney, E., Thompson, A., Comeault, A.A., Peede, D., D'Agostino, E.R., Pelaez, J. and Aguilar, J.M., 2021. Highly contiguous assemblies of 101 drosophilid genomes. *Elife*, *10*, p.e66405. |
| D. ambigua - D. pseudoobscura | *D.pseudoobscura* | SAMN16729702 | Kim, B.Y., Wang, J.R., Miller, D.E., Barmina, O., Delaney, E., Thompson, A., Comeault, A.A., Peede, D., D'Agostino, E.R., Pelaez, J. and Aguilar, J.M., 2021. Highly contiguous assemblies of 101 drosophilid genomes. *Elife*, *10*, p.e66405. |
| D. americana - D. lacicola | *D. lacicola* | SAMN29503225 | Yusuf, L.H., Tyukmaeva, V., Hoikkala, A. and Ritchie, M.G., 2022. Divergence and introgression among the virilis group of Drosophila. *Evolution Letters*, *6*(6), pp.537-551. |

| | | | |
|---|---|---|---|
| D. americana - D. littoralis | *D. littoralis* | SAMN29503227 | Yusuf, L.H., Tyukmaeva, V., Hoikkala, A. and Ritchie, M.G., 2022. Divergence and introgression among the virilis group of Drosophila. *Evolution Letters*, *6*(6), pp.537-551. |
| D. americana - D. montana | *D. montana* | SAMN29503231 | Yusuf, L.H., Tyukmaeva, V., Hoikkala, A. and Ritchie, M.G., 2022. Divergence and introgression among the virilis group of Drosophila. *Evolution Letters*, *6*(6), pp.537-551. |
| D. americana - D. virilis | *D. virilis* | SAMN16729705 | Kim, B.Y., Wang, J.R., Miller, D.E., Barmina, O., Delaney, E., Thompson, A., Comeault, A.A., Peede, D., D'Agostino, E.R., Pelaez, J. and Aguilar, J.M., 2021. Highly contiguous assemblies of 101 drosophilid genomes. *Elife*, *10*, p.e66405. |
| D. arizonae - D. mojavensis | *D. mojavaensis* | SAMN16729700 | Kim, B.Y., Wang, J.R., Miller, D.E., Barmina, O., Delaney, E., Thompson, A., Comeault, A.A., Peede, D., D'Agostino, E.R., Pelaez, J. and Aguilar, J.M., 2021. Highly contiguous assemblies of 101 drosophilid genomes. *Elife*, *10*, p.e66405. |
| D. auraria- D. rufa | *D. auraria* | SAMN12263403 | Bronski, M.J., Martinez, C.C., Weld, H.A. and Eisen, M.B., 2020. Whole genome sequences of 23 species from the Drosophila montium species group (Diptera: Drosophilidae): a resource for testing evolutionary hypotheses. *G3: Genes, Genomes, Genetics*, *10*(5), pp.1443-1455. |
| D. bipectinata - D. malerkotliana | *D. bipectinata* | SAMN16729609 | Kim, B.Y., Wang, J.R., Miller, D.E., Barmina, O., Delaney, E., Thompson, A., Comeault, A.A., Peede, D., D'Agostino, E.R., Pelaez, J. and Aguilar, J.M., 2021. Highly contiguous assemblies of 101 drosophilid genomes. *Elife*, *10*, p.e66405. |
| D. bipectinata - D. parabipectinata | *D. bipectinata* | SAMN16729609 | Kim, B.Y., Wang, J.R., Miller, D.E., Barmina, O., Delaney, E., Thompson, A., Comeault, A.A., Peede, D., D'Agostino, E.R., Pelaez, J. and Aguilar, J.M., 2021. Highly contiguous assemblies of 101 drosophilid genomes. *Elife*, *10*, p.e66405. |

| | | | |
|---|---|---|---|
| D. bipectinata - D. pseudoananassae | *D. bipectinata* | SAMN16729609 | Kim, B.Y., Wang, J.R., Miller, D.E., Barmina, O., Delaney, E., Thompson, A., Comeault, A.A., Peede, D., D'Agostino, E.R., Pelaez, J. and Aguilar, J.M., 2021. Highly contiguous assemblies of 101 drosophilid genomes. *Elife*, *10*, p.e66405. |
| D. borealis - D. lacicola | *D. lacicola* | SAMN29503225 | Yusuf, L.H., Tyukmaeva, V., Hoikkala, A. and Ritchie, M.G., 2022. Divergence and introgression among the virilis group of Drosophila. *Evolution Letters*, *6*(6), pp.537-551. |
| D. borealis - D. littoralis | *D. littoralis* | SAMN29503227 | Yusuf, L.H., Tyukmaeva, V., Hoikkala, A. and Ritchie, M.G., 2022. Divergence and introgression among the virilis group of Drosophila. *Evolution Letters*, *6*(6), pp.537-551. |
| D. borealis - D. montana | *D. montana* | SAMN29503231 | Yusuf, L.H., Tyukmaeva, V., Hoikkala, A. and Ritchie, M.G., 2022. Divergence and introgression among the virilis group of Drosophila. *Evolution Letters*, *6*(6), pp.537-551. |
| D. erecta - D. orena | *D. erecta* | SAMN16729697 | Kim, B.Y., Wang, J.R., Miller, D.E., Barmina, O., Delaney, E., Thompson, A., Comeault, A.A., Peede, D., D'Agostino, E.R., Pelaez, J. and Aguilar, J.M., 2021. Highly contiguous assemblies of 101 drosophilid genomes. *Elife*, *10*, p.e66405. |
| D. flavomontana - D. montana | *D. montana* | SAMN29503231 | Yusuf, L.H., Tyukmaeva, V., Hoikkala, A. and Ritchie, M.G., 2022. Divergence and introgression among the virilis group of Drosophila. *Evolution Letters*, *6*(6), pp.537-551. |
| D. insularis - D. willistoni | *D. willistoni* | SAMN16729706 | Kim, B.Y., Wang, J.R., Miller, D.E., Barmina, O., Delaney, E., Thompson, A., Comeault, A.A., Peede, D., D'Agostino, E.R., Pelaez, J. and Aguilar, J.M., 2021. Highly contiguous assemblies of 101 drosophilid genomes. *Elife*, *10*, p.e66405. |

| | | | |
|---|---|---|---|
| D. jambulina - D. punjabiensis | *D. punjabiensis* | SAMN12263408 | Bronski, M.J., Martinez, C.C., Weld, H.A. and Eisen, M.B., 2020. Whole genome sequences of 23 species from the Drosophila montium species group (Diptera: Drosophilidae): a resource for testing evolutionary hypotheses. *G3: Genes, Genomes, Genetics*, *10*(5), pp.1443-1455. |
| D. kanekoi - D. ezoana | *D. ezoana* | SAMN29503221 | Yusuf, L.H., Tyukmaeva, V., Hoikkala, A. and Ritchie, M.G., 2022. Divergence and introgression among the virilis group of Drosophila. *Evolution Letters*, *6*(6), pp.537-551. |
| D. kanekoi - D. littoralis | *D. littoralis* | SAMN29503227 | Yusuf, L.H., Tyukmaeva, V., Hoikkala, A. and Ritchie, M.G., 2022. Divergence and introgression among the virilis group of Drosophila. *Evolution Letters*, *6*(6), pp.537-551. |
| D. kanekoi - D. lummei | *D. lummei* | SAMN29503229 | Yusuf, L.H., Tyukmaeva, V., Hoikkala, A. and Ritchie, M.G., 2022. Divergence and introgression among the virilis group of Drosophila. *Evolution Letters*, *6*(6), pp.537-551. |
| D. kanekoi - D. virilis | *D. virilis* | SAMN16729705 | Kim, B.Y., Wang, J.R., Miller, D.E., Barmina, O., Delaney, E., Thompson, A., Comeault, A.A., Peede, D., D'Agostino, E.R., Pelaez, J. and Aguilar, J.M., 2021. Highly contiguous assemblies of 101 drosophilid genomes. *Elife*, *10*, p.e66405. |
| D. kepulauna - D. albomicans | *D. albomicans* | SAMN12703415 (legacy) | Mai, D., Nalley, M.J. and Bachtrog, D., 2020. Patterns of genomic differentiation in the Drosophila nasuta species complex. *Molecular biology and evolution*, *37*(1), pp.208-220; Wei, K.H.C., Mai, D., Chatla, K. and Bachtrog, D., 2022. Dynamics and impacts of transposable element proliferation in the Drosophila nasuta species group radiation. Molecular Biology and Evolution, 39(5), p.msac080. |

| | | | |
|---|---|---|---|
| D. kikkawai- D. leontia | *D. kikkawai* | SAMN16729629 | Kim, B.Y., Wang, J.R., Miller, D.E., Barmina, O., Delaney, E., Thompson, A., Comeault, A.A., Peede, D., D'Agostino, E.R., Pelaez, J. and Aguilar, J.M., 2021. Highly contiguous assemblies of 101 drosophilid genomes. *Elife*, *10*, p.e66405; Suvorov, A., Kim, B.Y., Wang, J., Armstrong, E.E., Peede, D., D'agostino, E.R., Price, D.K., Waddell, P.J., Lang, M., Courtier-Orgogozo, V. and David, J.R., 2022. Widespread introgression across a phylogeny of 155 Drosophila genomes. Current Biology, 32(1), pp.111-123. |
| D. kohkoa - D. albomicans | *D. albomicans* | SAMN12703415 (legacy) | Mai, D., Nalley, M.J. and Bachtrog, D., 2020. Patterns of genomic differentiation in the Drosophila nasuta species complex. *Molecular biology and evolution*, *37*(1), pp.208-220; Wei, K.H.C., Mai, D., Chatla, K. and Bachtrog, D., 2022. Dynamics and impacts of transposable element proliferation in the Drosophila nasuta species group radiation. Molecular Biology and Evolution, 39(5), p.msac080. |
| D. littoralis - D. ezoana | *D. ezoana* | SAMN29503221 | Yusuf, L.H., Tyukmaeva, V., Hoikkala, A. and Ritchie, M.G., 2022. Divergence and introgression among the virilis group of Drosophila. *Evolution Letters*, *6*(6), pp.537-551. |
| D. littoralis - D. montana | *D. montana* | SAMN29503231 | Yusuf, L.H., Tyukmaeva, V., Hoikkala, A. and Ritchie, M.G., 2022. Divergence and introgression among the virilis group of Drosophila. *Evolution Letters*, *6*(6), pp.537-551. |
| D. loweii - D. pseudoobscura | *D. pseudoobscura* | SAMN16729702 | Kim, B.Y., Wang, J.R., Miller, D.E., Barmina, O., Delaney, E., Thompson, A., Comeault, A.A., Peede, D., D'Agostino, E.R., Pelaez, J. and Aguilar, J.M., 2021. Highly contiguous assemblies of 101 drosophilid genomes. *Elife*, *10*, p.e66405. |

| | | | |
|---|---|---|---|
| D. lummei - D. ezoana | *D. lummei* | SAMN29503229 | Yusuf, L.H., Tyukmaeva, V., Hoikkala, A. and Ritchie, M.G., 2022. Divergence and introgression among the virilis group of Drosophila. *Evolution Letters*, *6*(6), pp.537-551. |
| D. lummei - D. littoralis | *D. lummei* | SAMN29503229 | Yusuf, L.H., Tyukmaeva, V., Hoikkala, A. and Ritchie, M.G., 2022. Divergence and introgression among the virilis group of Drosophila. *Evolution Letters*, *6*(6), pp.537-551. |
| D. lummei - D. virilis | *D. lummei* | SAMN29503229 | Yusuf, L.H., Tyukmaeva, V., Hoikkala, A. and Ritchie, M.G., 2022. Divergence and introgression among the virilis group of Drosophila. *Evolution Letters*, *6*(6), pp.537-551. |
| D. malerkotliana - D. parabipectinata | *D. parabipectinata* | SAMN16729611 | Kim, B.Y., Wang, J.R., Miller, D.E., Barmina, O., Delaney, E., Thompson, A., Comeault, A.A., Peede, D., D'Agostino, E.R., Pelaez, J. and Aguilar, J.M., 2021. Highly contiguous assemblies of 101 drosophilid genomes. *Elife*, *10*, p.e66405. |
| D. malerkotliana - D. pseudoananassae | *D. malerkotliana* | SAMN16729617 | Kim, B.Y., Wang, J.R., Miller, D.E., Barmina, O., Delaney, E., Thompson, A., Comeault, A.A., Peede, D., D'Agostino, E.R., Pelaez, J. and Aguilar, J.M., 2021. Highly contiguous assemblies of 101 drosophilid genomes. *Elife*, *10*, p.e66405. |
| D. mauritiana - D. erecta | *D. erecta* | SAMN16729697 | Kim, B.Y., Wang, J.R., Miller, D.E., Barmina, O., Delaney, E., Thompson, A., Comeault, A.A., Peede, D., D'Agostino, E.R., Pelaez, J. and Aguilar, J.M., 2021. Highly contiguous assemblies of 101 drosophilid genomes. *Elife*, *10*, p.e66405. |
| D. mauritiana - D. orena | *D. mauritiana* | SAMN06827991 | No specific publication; see https://www.ncbi.nlm.nih.gov/datasets/genome/GCA_004382145.1/ |
| D. mauritiana - D. sechellia | *D. mauritiana* | SAMN06827991 | No specific publication; see https://www.ncbi.nlm.nih.gov/datasets/genome/GCA_004382145.1/ |

| | | | |
|---|---|---|---|
| D. mauritiana - D. teissieri | *D. mauritiana* | SAMN06827991 | No specific publication; see https://www.ncbi.nlm.nih.gov/datasets/genome/GCA_004382145.1/ |
| D. mauritiana - D. yakuba | *D. yakuba* | SAMN16729707 | Kim, B.Y., Wang, J.R., Miller, D.E., Barmina, O., Delaney, E., Thompson, A., Comeault, A.A., Peede, D., D'Agostino, E.R., Pelaez, J. and Aguilar, J.M., 2021. Highly contiguous assemblies of 101 drosophilid genomes. *Elife*, *10*, p.e66405. |
| D. melanogaster - D. erecta | *D. erecta* | SAMN16729697 | Kim, B.Y., Wang, J.R., Miller, D.E., Barmina, O., Delaney, E., Thompson, A., Comeault, A.A., Peede, D., D'Agostino, E.R., Pelaez, J. and Aguilar, J.M., 2021. Highly contiguous assemblies of 101 drosophilid genomes. *Elife*, *10*, p.e66405. |
| D. melanogaster - D. orena | *D. melanogaster* | dmel_r6.39_FB2021_02/ | Flybase; v6.39. |
| D. melanogaster - D. sechellia | *D. melanogaster* | dmel_r6.39_FB2021_02/ | Flybase; v6.39. |
| D. melanogaster - D. simulans | *D. simulans* | SAMN16729704 | Kim, B.Y., Wang, J.R., Miller, D.E., Barmina, O., Delaney, E., Thompson, A., Comeault, A.A., Peede, D., D'Agostino, E.R., Pelaez, J. and Aguilar, J.M., 2021. Highly contiguous assemblies of 101 drosophilid genomes. *Elife*, *10*, p.e66405. |
| D. melanogaster - D. teissieri | *D. teissieri* | SAMN16729662 | Kim, B.Y., Wang, J.R., Miller, D.E., Barmina, O., Delaney, E., Thompson, A., Comeault, A.A., Peede, D., D'Agostino, E.R., Pelaez, J. and Aguilar, J.M., 2021. Highly contiguous assemblies of 101 drosophilid genomes. *Elife*, *10*, p.e66405; Suvorov, A., Kim, B.Y., Wang, J., Armstrong, E.E., Peede, D., D’agostino, E.R., Price, D.K., Waddell, P.J., Lang, M., Courtier-Orgogozo, V. and David, J.R., 2022. Widespread introgression across a phylogeny of 155 Drosophila genomes. Current Biology, 32(1), pp.111-123. |

| | | | |
|---|---|---|---|
| D. melanogaster - D. yakuba | *D. yakuba* | SAMN16729707 | Kim, B.Y., Wang, J.R., Miller, D.E., Barmina, O., Delaney, E., Thompson, A., Comeault, A.A., Peede, D., D'Agostino, E.R., Pelaez, J. and Aguilar, J.M., 2021. Highly contiguous assemblies of 101 drosophilid genomes. *Elife*, *10*, p.e66405. |
| D. miranda - D. persimilis | *D. persimilis* | SAMN16729701 | Kim, B.Y., Wang, J.R., Miller, D.E., Barmina, O., Delaney, E., Thompson, A., Comeault, A.A., Peede, D., D'Agostino, E.R., Pelaez, J. and Aguilar, J.M., 2021. Highly contiguous assemblies of 101 drosophilid genomes. *Elife*, *10*, p.e66405. |
| D. miranda - D. pseudoobscura | *D.pseudoobscura* | SAMN16729702 | Kim, B.Y., Wang, J.R., Miller, D.E., Barmina, O., Delaney, E., Thompson, A., Comeault, A.A., Peede, D., D'Agostino, E.R., Pelaez, J. and Aguilar, J.M., 2021. Highly contiguous assemblies of 101 drosophilid genomes. *Elife*, *10*, p.e66405. |
| D. nasuta - D. albomicans | *D. albomicans* | SAMN12703415 (legacy) | Mai, D., Nalley, M.J. and Bachtrog, D., 2020. Patterns of genomic differentiation in the Drosophila nasuta species complex. *Molecular biology and evolution*, *37*(1), pp.208-220; Wei, K.H.C., Mai, D., Chatla, K. and Bachtrog, D., 2022. Dynamics and impacts of transposable element proliferation in the Drosophila nasuta species group radiation. Molecular Biology and Evolution, 39(5), p.msac080. |
| D. novamexicana - D. lacicola | *D. novamexicana* | SAMN09383009 | No specific publication; see https://www.ncbi.nlm.nih.gov/datasets/genome/GCF_003285875.2/ |
| D. novamexicana - D. littoralis | *D. novamexicana* | SAMN09383009 | No specific publication; see https://www.ncbi.nlm.nih.gov/datasets/genome/GCF_003285875.2/ |
| D. novamexicana - D. montana | *D. novamexicana* | SAMN09383009 | No specific publication; see https://www.ncbi.nlm.nih.gov/datasets/genome/GCF_003285875.2/ |

| | | | |
|---|---|---|---|
| D. novamexicana - D. virilis | *D. novamexicana* | SAMN09383009 | No specific publication; see https://www.ncbi.nlm.nih.gov/datasets/genome/GCF_003285875.2/ |
| D. obscura - D. tristis | *D. obscura* | SAMN16729671 | Kim, B.Y., Wang, J.R., Miller, D.E., Barmina, O., Delaney, E., Thompson, A., Comeault, A.A., Peede, D., D'Agostino, E.R., Pelaez, J. and Aguilar, J.M., 2021. Highly contiguous assemblies of 101 drosophilid genomes. *Elife*, *10*, p.e66405. |
| D. parabipectinata - D. pseudoananassae | *D. parabipectinata* | SAMN16729611 | Kim, B.Y., Wang, J.R., Miller, D.E., Barmina, O., Delaney, E., Thompson, A., Comeault, A.A., Peede, D., D'Agostino, E.R., Pelaez, J. and Aguilar, J.M., 2021. Highly contiguous assemblies of 101 drosophilid genomes. *Elife*, *10*, p.e66405. |
| D. persimilis - D. bifasciata | *D. persimilis* | SAMN16729701 | Kim, B.Y., Wang, J.R., Miller, D.E., Barmina, O., Delaney, E., Thompson, A., Comeault, A.A., Peede, D., D'Agostino, E.R., Pelaez, J. and Aguilar, J.M., 2021. Highly contiguous assemblies of 101 drosophilid genomes. *Elife*, *10*, p.e66405. |
| D. persimilis - D. pseudoobscura | *D. persimilis* | SAMN16729701 | Kim, B.Y., Wang, J.R., Miller, D.E., Barmina, O., Delaney, E., Thompson, A., Comeault, A.A., Peede, D., D'Agostino, E.R., Pelaez, J. and Aguilar, J.M., 2021. Highly contiguous assemblies of 101 drosophilid genomes. *Elife*, *10*, p.e66405. |
| D. persimilis - D. subobscura | *D. persimilis* | SAMN16729701 | Kim, B.Y., Wang, J.R., Miller, D.E., Barmina, O., Delaney, E., Thompson, A., Comeault, A.A., Peede, D., D'Agostino, E.R., Pelaez, J. and Aguilar, J.M., 2021. Highly contiguous assemblies of 101 drosophilid genomes. *Elife*, *10*, p.e66405. |
| D. pseudoobscura - D. bifasciata | *D.pseudoobscura* | SAMN16729702 | Kim, B.Y., Wang, J.R., Miller, D.E., Barmina, O., Delaney, E., Thompson, A., Comeault, A.A., Peede, D., D'Agostino, E.R., Pelaez, J. and Aguilar, J.M., 2021. Highly contiguous assemblies of 101 drosophilid genomes. *Elife*, *10*, p.e66405. |

| | | | |
|---|---|---|---|
| D. pseudoobscura - D. subobscura | *D.pseudoobscura* | SAMN16729702 | Kim, B.Y., Wang, J.R., Miller, D.E., Barmina, O., Delaney, E., Thompson, A., Comeault, A.A., Peede, D., D'Agostino, E.R., Pelaez, J. and Aguilar, J.M., 2021. Highly contiguous assemblies of 101 drosophilid genomes. *Elife*, *10*, p.e66405. |
| D. sechellia - D. erecta | *D. erecta* | SAMN16729697 | Kim, B.Y., Wang, J.R., Miller, D.E., Barmina, O., Delaney, E., Thompson, A., Comeault, A.A., Peede, D., D'Agostino, E.R., Pelaez, J. and Aguilar, J.M., 2021. Highly contiguous assemblies of 101 drosophilid genomes. *Elife*, *10*, p.e66405. |
| D. sechellia - D. orena | *D. melanogaster* | dmel_r6.39_FB2021_02/ | Flybase; v6.39. |
| D. sechellia - D. teissieri | *D. teissieri* | SAMN16729662 | Kim, B.Y., Wang, J.R., Miller, D.E., Barmina, O., Delaney, E., Thompson, A., Comeault, A.A., Peede, D., D'Agostino, E.R., Pelaez, J. and Aguilar, J.M., 2021. Highly contiguous assemblies of 101 drosophilid genomes. *Elife*, *10*, p.e66405; Suvorov, A., Kim, B.Y., Wang, J., Armstrong, E.E., Peede, D., D'agostino, E.R., Price, D.K., Waddell, P.J., Lang, M., Courtier-Orgogozo, V. and David, J.R., 2022. Widespread introgression across a phylogeny of 155 Drosophila genomes. Current Biology, 32(1), pp.111-123. |
| D. sechellia - D. yakuba | *D. yakuba* | SAMN16729707 | Kim, B.Y., Wang, J.R., Miller, D.E., Barmina, O., Delaney, E., Thompson, A., Comeault, A.A., Peede, D., D'Agostino, E.R., Pelaez, J. and Aguilar, J.M., 2021. Highly contiguous assemblies of 101 drosophilid genomes. *Elife*, *10*, p.e66405. |
| D. simulans - D. erecta | *D. simulans* | SAMN16729704 | Kim, B.Y., Wang, J.R., Miller, D.E., Barmina, O., Delaney, E., Thompson, A., Comeault, A.A., Peede, D., D'Agostino, E.R., Pelaez, J. and Aguilar, J.M., 2021. Highly contiguous assemblies of 101 drosophilid genomes. *Elife*, *10*, p.e66405. |

| | | | |
|---|---|---|---|
| D. simulans - D. orena | *D. simulans* | SAMN16729704 | Kim, B.Y., Wang, J.R., Miller, D.E., Barmina, O., Delaney, E., Thompson, A., Comeault, A.A., Peede, D., D'Agostino, E.R., Pelaez, J. and Aguilar, J.M., 2021. Highly contiguous assemblies of 101 drosophilid genomes. *Elife*, *10*, p.e66405. |
| D. simulans - D. teissieri | *D. simulans* | SAMN16729704 | Kim, B.Y., Wang, J.R., Miller, D.E., Barmina, O., Delaney, E., Thompson, A., Comeault, A.A., Peede, D., D'Agostino, E.R., Pelaez, J. and Aguilar, J.M., 2021. Highly contiguous assemblies of 101 drosophilid genomes. *Elife*, *10*, p.e66405. |
| D. simulans - D. yakuba | *D. simulans* | SAMN16729704 | Kim, B.Y., Wang, J.R., Miller, D.E., Barmina, O., Delaney, E., Thompson, A., Comeault, A.A., Peede, D., D'Agostino, E.R., Pelaez, J. and Aguilar, J.M., 2021. Highly contiguous assemblies of 101 drosophilid genomes. *Elife*, *10*, p.e66405. |
| D. subobscura - D. ambigua | *D. subobscura* | SAMN16729670 | Kim, B.Y., Wang, J.R., Miller, D.E., Barmina, O., Delaney, E., Thompson, A., Comeault, A.A., Peede, D., D'Agostino, E.R., Pelaez, J. and Aguilar, J.M., 2021. Highly contiguous assemblies of 101 drosophilid genomes. *Elife*, *10*, p.e66405. |
| D. subobscura - D. bifasciata | *D. subobscura* | SAMN16729670 | Kim, B.Y., Wang, J.R., Miller, D.E., Barmina, O., Delaney, E., Thompson, A., Comeault, A.A., Peede, D., D'Agostino, E.R., Pelaez, J. and Aguilar, J.M., 2021. Highly contiguous assemblies of 101 drosophilid genomes. *Elife*, *10*, p.e66405. |
| D. subobscura - D. obscura | *D. subobscura* | SAMN16729670 | Kim, B.Y., Wang, J.R., Miller, D.E., Barmina, O., Delaney, E., Thompson, A., Comeault, A.A., Peede, D., D'Agostino, E.R., Pelaez, J. and Aguilar, J.M., 2021. Highly contiguous assemblies of 101 drosophilid genomes. *Elife*, *10*, p.e66405. |

| | | | |
|---|---|---|---|
| D. subobscura - D. tristis | *D. subobscura* | SAMN16729670 | Kim, B.Y., Wang, J.R., Miller, D.E., Barmina, O., Delaney, E., Thompson, A., Comeault, A.A., Peede, D., D'Agostino, E.R., Pelaez, J. and Aguilar, J.M., 2021. Highly contiguous assemblies of 101 drosophilid genomes. *Elife*, *10*, p.e66405. |
| D. sulfurigaster albostrigata - D. kepulauna | *D. neonasuta* | SAMN06052360 | see https://www.ncbi.nlm.nih.gov/datasets/genome/GCA_005889595.1/ |
| D. sulfurigaster albostrigata - D. pulaua | *D. neonasuta* | SAMN06052360 | see https://www.ncbi.nlm.nih.gov/datasets/genome/GCA_005889595.1/ |
| D. sulfurigaster albostrigata - D. sulfurigaster bilimbata | *D. neonasuta* | SAMN06052360 | see https://www.ncbi.nlm.nih.gov/datasets/genome/GCA_005889595.1/ |
| D. sulfurigaster albostrigata - D. sulfurigaster neonasuta | *D. neonasuta* | SAMN06052360 | see https://www.ncbi.nlm.nih.gov/datasets/genome/GCA_005889595.1/ |
| D. sulfurigaster albostrigata - D. sulfurigaster sulfurigaster | *D. neonasuta* | SAMN06052360 | see https://www.ncbi.nlm.nih.gov/datasets/genome/GCA_005889595.1/ |
| D. sulfurigaster bilimbata - D. pulaua | *D. neonasuta* | SAMN06052360 | see https://www.ncbi.nlm.nih.gov/datasets/genome/GCA_005889595.1/ |

| | | | |
|---|---|---|---|
| D. sulfurigaster bilimbata - D. sulfurigaster neonasuta | *D. neonasuta* | SAMN06052360 | see https://www.ncbi.nlm.nih.gov/datasets/genome/GCA_005889595.1/ |
| D. sulfurigaster bilimbata - D. sulfurigaster sulfurigaster | *D. neonasuta* | SAMN06052360 | see https://www.ncbi.nlm.nih.gov/datasets/genome/GCA_005889595.1/ |
| D. sulfurigaster neonasuta - D. pulaua | *D. neonasuta* | SAMN06052360 | see https://www.ncbi.nlm.nih.gov/datasets/genome/GCA_005889595.1/ |
| D. sulfurigaster neonasuta - D. sulfurigaster sulfurigaster | *D. neonasuta* | SAMN06052360 | see https://www.ncbi.nlm.nih.gov/datasets/genome/GCA_005889595.1/ |
| D. sulfurigaster sulfurigaster - D. pulaua | *D. neonasuta* | SAMN06052360 | see https://www.ncbi.nlm.nih.gov/datasets/genome/GCA_005889595.1/ |
| D. teissieri-D. erecta | *D. teissieri* | SAMN16729662 | Kim, B.Y., Wang, J.R., Miller, D.E., Barmina, O., Delaney, E., Thompson, A., Comeault, A.A., Peede, D., D'Agostino, E.R., Pelaez, J. and Aguilar, J.M., 2021. Highly contiguous assemblies of 101 drosophilid genomes. *Elife*, *10*, p.e66405; Suvorov, A., Kim, B.Y., Wang, J., Armstrong, E.E., Peede, D., D'agostino, E.R., Price, D.K., Waddell, P.J., Lang, M., Courtier-Orgogozo, V. and David, J.R., 2022. Widespread introgression across a phylogeny of 155 Drosophila genomes. Current Biology, 32(1), pp.111-123. |

| | | | |
|---|---|---|---|
| D. teissieri - D. orena | *D. teissieri* | SAMN16729662 | Kim, B.Y., Wang, J.R., Miller, D.E., Barmina, O., Delaney, E., Thompson, A., Comeault, A.A., Peede, D., D'Agostino, E.R., Pelaez, J. and Aguilar, J.M., 2021. Highly contiguous assemblies of 101 drosophilid genomes. *Elife*, *10*, p.e66405; Suvorov, A., Kim, B.Y., Wang, J., Armstrong, E.E., Peede, D., D'agostino, E.R., Price, D.K., Waddell, P.J., Lang, M., Courtier-Orgogozo, V. and David, J.R., 2022. Widespread introgression across a phylogeny of 155 Drosophila genomes. Current Biology, 32(1), pp.111-123. |
| D. triauraria - D. rufa | *D. triauraria* | SAMN16744625 | Kim, B.Y., Wang, J.R., Miller, D.E., Barmina, O., Delaney, E., Thompson, A., Comeault, A.A., Peede, D., D'Agostino, E.R., Pelaez, J. and Aguilar, J.M., 2021. Highly contiguous assemblies of 101 drosophilid genomes. *Elife*, *10*, p.e66405. |
| D. virilis - D. ezoana | *D. ezoana* | SAMN29503221 | Yusuf, L.H., Tyukmaeva, V., Hoikkala, A. and Ritchie, M.G., 2022. Divergence and introgression among the virilis group of Drosophila. *Evolution Letters*, *6*(6), pp.537-551. |
| D. virilis - D. lacicola | *D. lacicola* | SAMN29503225 | Yusuf, L.H., Tyukmaeva, V., Hoikkala, A. and Ritchie, M.G., 2022. Divergence and introgression among the virilis group of Drosophila. *Evolution Letters*, *6*(6), pp.537-551. |
| D. virilis - D. littoralis | *D. virilis* | SAMN16729705 | Kim, B.Y., Wang, J.R., Miller, D.E., Barmina, O., Delaney, E., Thompson, A., Comeault, A.A., Peede, D., D'Agostino, E.R., Pelaez, J. and Aguilar, J.M., 2021. Highly contiguous assemblies of 101 drosophilid genomes. *Elife*, *10*, p.e66405. |
| D. yakuba - D. erecta | *D. yakuba* | SAMN16729707 | Kim, B.Y., Wang, J.R., Miller, D.E., Barmina, O., Delaney, E., Thompson, A., Comeault, A.A., Peede, D., D'Agostino, E.R., Pelaez, J. and Aguilar, J.M., 2021. Highly contiguous assemblies of 101 drosophilid genomes. *Elife*, *10*, p.e66405. |

| | | | |
|---|---|---|---|
| D. yakuba - D. orena | *D. yakuba* | SAMN16729707 | Kim, B.Y., Wang, J.R., Miller, D.E., Barmina, O., Delaney, E., Thompson, A., Comeault, A.A., Peede, D., D'Agostino, E.R., Pelaez, J. and Aguilar, J.M., 2021. Highly contiguous assemblies of 101 drosophilid genomes. *Elife*, *10*, p.e66405. |
| D. yakuba - D. santomea | *D. yakuba* | SAMN16729707 | Kim, B.Y., Wang, J.R., Miller, D.E., Barmina, O., Delaney, E., Thompson, A., Comeault, A.A., Peede, D., D'Agostino, E.R., Pelaez, J. and Aguilar, J.M., 2021. Highly contiguous assemblies of 101 drosophilid genomes. *Elife*, *10*, p.e66405. |
| D. yakuba - D. teissieri | *D. yakuba* | SAMN16729707 | Kim, B.Y., Wang, J.R., Miller, D.E., Barmina, O., Delaney, E., Thompson, A., Comeault, A.A., Peede, D., D'Agostino, E.R., Pelaez, J. and Aguilar, J.M., 2021. Highly contiguous assemblies of 101 drosophilid genomes. *Elife*, *10*, p.e66405. |

**Supplementary Table 3:** Species pairs and the genomes used to map whole-genome sequences to. Includes species' genome which was used for analysis, NCBI BioSample identification of the genome used for each pair and the publications where the data sequenced and assembled (where possible).

| Pairs | Sample | Coverage (mean) | Coverage (SD) | Coverage (min) | Coverage (max) |
|---|---|---|---|---|---|
| *D. affinis - D. athabasca* | SRR9967640 | 13.35 | 7.86 | 2 | 29 |
| *D. affinis - D. athabasca* | ERR127384 | 16.24 | 16.86 | 2 | 49 |
| *D. ambigua - D. obscura* | SRR13070667 | 18.55 | 15.92 | 2 | 50 |
| *D. ambigua - D. obscura* | SRR13070671 | 16.3 | 10.81 | 2 | 37 |
| *D. ambigua - D. persimilis* | SRR12899691 | 24.45 | 19.76 | 2 | 63 |
| *D. ambigua - D. persimilis* | SRR13070667 | 16.89 | 20.92 | 2 | 58 |
| *D. ambigua - D. pseudoobscura* | SRR11230572 | 25.82 | 14.29 | 2 | 54 |
| *D. ambigua - D. pseudoobscura* | SRR13070667 | 16.93 | 22.35 | 2 | 61 |
| *D. ambigua - D. pseudoobscura* | SRR13070669 | 15.01 | 30.76 | 2 | 76 |
| *D. ambigua - D. pseudoobscura* | SRR13070667 | 14.68 | 25.11 | 2 | 64 |
| *D. americana - D. lacicola* | ERR2610695 | 16.82 | 44.13 | 2 | 105 |
| *D. americana - D. lacicola* | SRR20226091 | 22 | 8.95 | 2 | 39 |
| *D. americana - D. littoralis* | ERR2610695 | 16.71 | 22.26 | 2 | 61 |
| *D. americana - D. littoralis* | SRR13070670 | 16.28 | 33.6 | 2 | 83 |
| *D. americana - D. montana* | SRR3218635 | 28.99 | 56.14 | 2 | 141 |
| *D. americana - D. montana* | ERR2610695 | 16.66 | 209.99 | 2 | 436 |
| *D. americana - D. novamexicana* | ERR2610692 | 19.69 | 61.5 | 2 | 142 |
| *D. americana - D. novamexicana* | ERR2610695 | 19.83 | 48.49 | 2 | 116 |
| *D. americana - D. virilis* | SRR9426114 | 9.94 | 8.52 | 2 | 26 |
| *D. americana - D. virilis* | ERR2610695 | 18.23 | 47.92 | 2 | 114 |
| *D. arizonae - D. mojavensis* | SRR2070760 | 10.93 | 10.28 | 2 | 31 |
| *D. arizonae - D. mojavensis* | SRR6425997 | 30.01 | 15.74 | 2 | 61 |
| *D. auraria - D. rufa* | SRR13188937 | 11.34 | 10.05 | 2 | 31 |
| *D. auraria - D. rufa* | SRR13070713 | 14.91 | 16.77 | 2 | 48 |
| *D. auraria - D. triauraria* | SRR13188937 | 11.65 | 9.47 | 2 | 30 |
| *D. auraria - D. triauraria* | SRR13188922 | 16.05 | 12.22 | 2 | 40 |

| | | | | | |
|---|---|---|---|---|---|
| *D. bipectinata - D. malerkotliana (pallens)* | SRR15048573 | 9.74 | 8.91 | 2 | 27 |
| *D. bipectinata - D. malerkotliana (pallens)* | SRR13070716 | 19.17 | 14.54 | 2 | 48 |
| *D. bipectinata - D. parabipectinata* | SRR13070721 | 12.33 | 12.51 | 2 | 37 |
| *D. bipectinata - D. parabipectinata* | SRR15048573 | 9.74 | 8.91 | 2 | 27 |
| *D. bipectinata - D. pseudoananassae (nigrens)* | SRR15048573 | 9.74 | 8.91 | 2 | 27 |
| *D. bipectinata - D. pseudoananassae (nigrens)* | SRR13070715 | 11.19 | 17.17 | 2 | 45 |
| *D. borealis - D. lacicola* | SRR13070670 | 21.49 | 18.57 | 2 | 58 |
| *D. borealis - D. lacicola* | SRR20226091 | 22 | 8.95 | 2 | 39 |
| *D. borealis - D. littoralis* | SRR13070670 | 16.28 | 33.6 | 2 | 83 |
| *D. borealis - D. littoralis* | SRR13070670 | 20.35 | 17.55 | 2 | 55 |
| *D. borealis - D. montana* | SRR3218635 | 28.99 | 56.13 | 2 | 141 |
| *D. borealis - D. montana* | SRR13070670 | 20.76 | 14.82 | 2 | 50 |
| *D. erecta - D. orena* | SRR1977592 | 9.8 | 8.49 | 2 | 26 |
| *D. erecta - D. orena* | SRR6425990 | 40.14 | 17.62 | 2 | 75 |
| *D. flavomontana - D. montana* | SRR3218635 | 28.99 | 56.15 | 2 | 141 |
| *D. flavomontana - D. montana* | SRR20226092 | 21.67 | 87.7 | 2 | 197 |
| *D. heteroneura - D. planitibia* | SRR2233094 | 17.54 | 12.91 | 2 | 43 |
| *D. heteroneura - D. planitibia* | SRR2233332 | 29.52 | 202.99 | 2 | 435 |
| *D. heteroneura - D. silvestris* | SRR2233094 | 17.54 | 12.91 | 2 | 43 |
| *D. heteroneura - D. silvestris* | SRR2233314 | 18.35 | 12.57 | 2 | 43 |
| *D. insularis - D. willistoni* | SRR6426003 | 23.82 | 19.33 | 2 | 62 |
| *D. insularis - D. willistoni* | SRR13070692 | 31.88 | 37.87 | 2 | 107 |
| *D. jambulina - D. punjabiensis* | SRR9997923 | 18.72 | 9.44 | 2 | 37 |
| *D. jambulina - D. punjabiensis* | SRR15048354 | 7 | 23.94 | 2 | 54 |
| *D. kanekoi - D. ezoana* | SRR20226095 | 22.75 | 10.2 | 2 | 43 |

| | | | | | |
|---|---|---|---|---|---|
| *D. kanekoi - D. ezoana* | SRR20226093 | 19.5 | 461.55 | 2 | 942 |
| *D. kanekoi - D. littoralis* | SRR13070670 | 16.28 | 33.6 | 2 | 83 |
| *D. kanekoi - D. littoralis* | SRR20226093 | 20.22 | 24.8 | 2 | 69 |
| *D. kanekoi - D. virilis* | SRR20226093 | 19.6 | 282.26 | 2 | 584 |
| *D. kanekoi - D. virilis* | SRR5278982 | 16.68 | 12.33 | 2 | 41 |
| *kanekoi_virilis* | SRR9426114 | 9.94 | 8.52 | 2 | 26 |
| *kanekoi_virilis* | SRR20226093 | 20.62 | 12.68 | 2 | 45 |
| *D. kepulauna - D. albomicans* | DRR076000 | 82.24 | 454.35 | 2 | 990 |
| *D. kepulauna - D. albomicans* | SRR10100373 | 9.19 | 5.66 | 2 | 20 |
| *D. kikkawai - D. leontia* | SRR1810494 | 9.04 | 8.68 | 2 | 26 |
| *D. kikkawai - D. leontia* | SRR15048447 | 9.23 | 10.13 | 2 | 29 |
| *D. kohkoa - D. albomicans* | DRR160734 | 49.97 | 68.1 | 2 | 186 |
| *D. kohkoa - D. albomicans* | DRR076000 | 82.24 | 454.36 | 2 | 990 |
| *D. lacicola - D. montana* | SRR3218635 | 28.99 | 56.14 | 2 | 141 |
| *D. lacicola - D. montana* | SRR20226091 | 20.65 | 10.58 | 2 | 41 |
| *D. littoralis - D. ezoana* | SRR20226095 | 22.75 | 10.2 | 2 | 43 |
| *D. littoralis - D. ezoana* | SRR13070670 | 15.09 | 58.14 | 2 | 131 |
| *D. littoralis - D. montana* | SRR13070670 | 15.21 | 36.84 | 2 | 88 |
| *D. littoralis - D. montana* | SRR3218635 | 28.99 | 56.13 | 2 | 141 |
| *D. loweii - D. pseudoobscura* | SRR11230572 | 25.82 | 14.29 | 2 | 54 |
| *D. loweii - D. pseudoobscura* | SRR9967663 | 12.31 | 18.12 | 2 | 48 |
| *D. lummei - D. ezoana* | SRR20226095 | 21.72 | 19.62 | 2 | 60 |
| *D. lummei - D. ezoana* | SRR5278982 | 16.68 | 12.33 | 2 | 41 |
| *D. lummei - D. littoralis* | SRR13070670 | 15.58 | 34.12 | 2 | 83 |
| *D. lummei - D. littoralis* | SRR5278982 | 16.68 | 12.33 | 2 | 41 |
| *D. lummei - D. virilis* | SRR9426114 | 8.86 | 8.21 | 2 | 25 |
| *D. lummei - D. virilis* | SRR5278982 | 16.68 | 12.33 | 2 | 41 |
| *D. malerkotliana (pallens) - D. parabipectinata* | SRR13070721 | 11.7 | 8.39 | 2 | 28 |

| | | | | | |
|---|---|---|---|---|---|
| *D. malerkotliana (pallens) - D. parabipectinata* | SRR13070716 | 18.9 | 13.71 | 2 | 46 |
| *D. malerkotliana (pallens) - D. pseudoananassae (nigrens)* | SRR13070716 | 19.54 | 11.85 | 2 | 43 |
| *D. malerkotliana (pallens) - D. pseudoananassae (nigrens)* | SRR13070715 | 11.3 | 15.78 | 2 | 42 |
| *D. mauritiana - D. erecta* | SRR1560109 | 12.82 | 8.14 | 2 | 29 |
| *D. mauritiana - D. erecta* | SRR6425990 | 40.14 | 17.62 | 2 | 75 |
| *D. mauritiana - D. orena* | SRR1560109 | 15.23 | 7.23 | 2 | 29 |
| *D. mauritiana - D. orena* | SRR1977592 | 8.85 | 7.3 | 2 | 23 |
| *D. mauritiana - D. sechellia* | SRR1560109 | 15.23 | 7.23 | 2 | 29 |
| *D. mauritiana - D. sechellia* | SRR14138507 | 22.38 | 13.35 | 2 | 49 |
| *D. mauritiana - D. teissieri* | SRR5860615 | 11.89 | 12.18 | 2 | 36 |
| *D. mauritiana - D. teissieri* | SRR1560109 | 15.23 | 7.23 | 2 | 29 |
| *D. mauritiana - D. yakuba* | SRR1560109 | 12.79 | 23.88 | 2 | 60 |
| *D. mauritiana - D. yakuba* | SRR9700025 | 16.48 | 7.73 | 2 | 31 |
| *D. melanogaster - D. erecta* | SRR3931586 | 13.19 | 132.76 | 2 | 278 |
| *D. melanogaster - D. erecta* | SRR6425990 | 40.14 | 17.62 | 2 | 75 |
| *D. melanogaster - D. orena* | SRR1977592 | 8.98 | 7.93 | 2 | 24 |
| *D. melanogaster - D. orena* | SRR3931586 | 12.21 | 12.3 | 2 | 36 |
| *D. melanogaster - D. sechellia* | SRR14138507 | 21.87 | 34.25 | 2 | 90 |
| *D. melanogaster - D. sechellia* | SRR3931586 | 12.21 | 12.3 | 2 | 36 |
| *D. melanogaster - D. simulans* | SRR11456787 | 26.86 | 12.34 | 2 | 51 |
| *D. melanogaster - D. simulans* | SRR3931586 | 13.15 | 13.95 | 2 | 41 |
| *D. melanogaster - D. teissieri* | SRR5860615 | 12.82 | 7.06 | 2 | 26 |
| *D. melanogaster - D. teissieri* | SRR3931586 | 12.83 | 89.37 | 2 | 191 |
| *D. melanogaster - D. yakuba* | SRR9700025 | 16.48 | 7.73 | 2 | 31 |
| *D. melanogaster - D. yakuba* | SRR3931586 | 12.84 | 78.35 | 2 | 169 |

| | | | | | |
|---|---|---|---|---|---|
| *D. miranda - D. bifasciata* | SRR789669 | 9.27 | 10.01 | 2 | 29 |
| *D. miranda - D. bifasciata* | DRR061004 | 46.87 | 24.65 | 2 | 96 |
| *D. miranda - D. persimilis* | SRR12899691 | 24.45 | 19.76 | 2 | 63 |
| *D. miranda - D. persimilis* | SRR789669 | 13.26 | 11 | 2 | 35 |
| *D. miranda - D. pseudoobscura* | SRR11230572 | 25.82 | 14.29 | 2 | 54 |
| *D. miranda - D. pseudoobscura* | SRR789669 | 13.23 | 11.16 | 2 | 35 |
| *D. miranda - D. subobscura* | SRR789669 | 9.3 | 8.52 | 2 | 26 |
| *D. miranda - D. subobscura* | SRR9967664 | 14.94 | 8.94 | 2 | 32 |
| *D. nasuta - D. albomicans* | SRR10100355 | 11.68 | 6.74 | 2 | 25 |
| *D. nasuta - D. albomicans* | DRR076000 | 82.24 | 454.36 | 2 | 990 |
| *D. novamexicana - D. lacicola* | SRR20226091 | 20.59 | 19.38 | 2 | 59 |
| *D. novamexicana - D. lacicola* | ERR2610692 | 19.7 | 61.49 | 2 | 142 |
| *D. novamexicana - D. littoralis* | ERR2610692 | 19.69 | 61.48 | 2 | 142 |
| *D. novamexicana - D. littoralis* | SRR13070670 | 16.35 | 20.22 | 2 | 56 |
| *D. novamexicana - D. montana* | SRR3218635 | 25.21 | 95.57 | 2 | 216 |
| *D. novamexicana - D. montana* | ERR2610692 | 19.7 | 61.51 | 2 | 142 |
| *D. novamexicana - D. virilis* | SRR9426114 | 9.32 | 8.61 | 2 | 26 |
| *D. novamexicana - D. virilis* | ERR2610692 | 19.7 | 61.48 | 2 | 142 |
| *D. obscura - D. tristis* | SRR13070671 | 16.3 | 10.81 | 2 | 37 |
| *D. obscura - D. tristis* | SRR13070669 | 19.21 | 18.36 | 2 | 55 |
| *D. parabipectinata - D. pseudoananassae* | SRR13070715 | 10.96 | 17.22 | 2 | 45 |
| *D. parabipectinata - D. pseudoananassae* | SRR13070721 | 11.7 | 8.39 | 2 | 28 |
| *D. persimilis - D. bifasciata* | SRR12899691 | 24.45 | 19.76 | 2 | 63 |
| *D. persimilis - D. bifasciata* | DRR061004 | 37.59 | 29.98 | 2 | 97 |
| *D. persimilis - D. pseudoobscura* | SRR11230572 | 25.43 | 15.99 | 2 | 57 |

| | | | | | |
|---|---|---|---|---|---|
| *D. persimilis - D. pseudoobscura* | SRR12899691 | 24.45 | 19.76 | 2 | 63 |
| *D. persimilis - D. subobscura* | SRR12899691 | 24.45 | 19.76 | 2 | 63 |
| *D. persimilis - D. subobscura* | SRR9967664 | 11.32 | 8.56 | 2 | 28 |
| *D. planitibia - D. silvestris* | SRR2233314 | 18.35 | 12.57 | 2 | 43 |
| *D. planitibia - D. silvestris* | SRR2233332 | 29.52 | 203.01 | 2 | 435 |
| *D. pseudoobscura - D. bifasciata* | SRR11230572 | 25.82 | 14.29 | 2 | 54 |
| *D. pseudoobscura - D. bifasciata* | DRR061004 | 37.7 | 28.83 | 2 | 95 |
| *D. pseudoobscura - D. subobscura* | SRR11230572 | 25.82 | 14.29 | 2 | 54 |
| *D. pseudoobscura - D. subobscura* | SRR9967664 | 11.35 | 9.13 | 2 | 29 |
| *D. sechellia - D. erecta* | SRR14138507 | 21.37 | 17.45 | 2 | 56 |
| *D. sechellia - D. erecta* | SRR6425990 | 40.14 | 17.62 | 2 | 75 |
| *D. sechellia - D. orena* | SRR1977592 | 8.98 | 7.93 | 2 | 24 |
| *D. sechellia - D. orena* | SRR14138507 | 21.87 | 34.25 | 2 | 90 |
| *D. sechellia - D. teissieri* | SRR5860615 | 12.82 | 7.06 | 2 | 26 |
| *D. sechellia - D. teissieri* | SRR14138507 | 21.06 | 17.34 | 2 | 55 |
| *D. sechellia - D. yakuba* | SRR14138507 | 21.14 | 23.78 | 2 | 68 |
| *D. sechellia - D. yakuba* | SRR9700025 | 16.48 | 7.73 | 2 | 31 |
| *D. simulans - D. erecta* | SRR6425990 | 34.9 | 19.93 | 2 | 74 |
| *D. simulans - D. erecta* | SRR11456787 | 26.86 | 12.34 | 2 | 51 |
| *D. simulans - D. orena* | SRR1977592 | 9.22 | 8.4 | 2 | 26 |
| *D. simulans - D. orena* | SRR11456787 | 26.86 | 12.34 | 2 | 51 |
| *D. simulans - D. sechellia* | SRR14138507 | 24.04 | 15.67 | 2 | 55 |
| *D. simulans - D. sechellia* | SRR11456787 | 26.86 | 12.34 | 2 | 51 |
| *D. simulans - D. teissieri* | SRR5860615 | 12.52 | 11.5 | 2 | 35 |
| *D. simulans - D. teissieri* | SRR11456787 | 26.86 | 12.34 | 2 | 51 |

| | | | | | |
|---|---|---|---|---|---|
| *D. simulans - D. yakuba* | SRR11456787 | 26.86 | 12.34 | 2 | 51 |
| *D. simulans - D. yakuba* | SRR9700025 | 12.97 | 8.79 | 2 | 30 |
| *D. subobscura - D. ambigua* | SRR9967664 | 14.94 | 8.94 | 2 | 32 |
| *D. subobscura - D. ambigua* | SRR13070667 | 16.97 | 25.77 | 2 | 68 |
| *D. subobscura - D. bifasciata* | SRR9967664 | 14.94 | 8.94 | 2 | 32 |
| *D. subobscura - D. bifasciata* | DRR061004 | 36.99 | 33.81 | 2 | 104 |
| *D. subobscura - D. obscura* | SRR13070671 | 14.36 | 31.75 | 2 | 77 |
| *D. subobscura - D. obscura* | SRR9967664 | 14.94 | 8.94 | 2 | 32 |
| *D. subobscura - D. tristis* | SRR9967664 | 14.94 | 8.94 | 2 | 32 |
| *D. subobscura - D. tristis* | SRR13070669 | 17.07 | 42.16 | 2 | 101 |
| *D. sulfurigaster albostrigata - D. kepulauna* | SRR10100407 | 5.58 | 6.02 | 2 | 17 |
| *D. sulfurigaster albostrigata - D. kepulauna* | SRR10100373 | 9.41 | 13.34 | 2 | 36 |
| *D. sulfurigaster albostrigata - D. pulaua* | SRR10100407 | 5.58 | 6.01 | 2 | 17 |
| *D. sulfurigaster albostrigata - D. pulaua* | SRR10100335 | 17.39 | 91.72 | 2 | 200 |
| *D. sulfurigaster albostrigata - D. sulfurigaster bilimbata* | SRR10100323 | 26.51 | 21.29 | 2 | 69 |
| *D. sulfurigaster albostrigata - D. sulfurigaster bilimbata* | SRR10100308 | 15.76 | 19.18 | 2 | 54 |
| *D. sulfurigaster albostrigata - D. sulfurigaster neonasuta* | SRR5059342 | 13.07 | 180.59 | 2 | 374 |
| *D. sulfurigaster albostrigata - D. sulfurigaster neonasuta* | SRR10100407 | 5.58 | 6.01 | 2 | 17 |
| *D. sulfurigaster albostrigata - D. sulfurigaster sulfurigaster* | SRR10100323 | 26.51 | 21.29 | 2 | 69 |
| *D. sulfurigaster albostrigata - D.* | SRR10100401 | 14.69 | 46.96 | 2 | 108 |

| | | | | | |
|---|---|---|---|---|---|
| *sulfurigaster sulfurigaster* | | | | | |
| *D. sulfurigaster bilimbata - pulaua* | SRR10100335 | 17.39 | 91.72 | 2 | 200 |
| *D. sulfurigaster bilimbata - pulaua* | SRR10100308 | 15.76 | 19.18 | 2 | 54 |
| *D. sulfurigaster bilimbata - sulfurigaster neonasuta* | SRR5059342 | 13.07 | 180.59 | 2 | 374 |
| *D. sulfurigaster bilimbata - sulfurigaster neonasuta* | SRR10100308 | 15.76 | 19.18 | 2 | 54 |
| *D. sulfurigaster bilimbata - D. sulfurigaster sulfurigaster* | SRR10100308 | 15.76 | 19.18 | 2 | 54 |
| *D. sulfurigaster bilimbata - D. sulfurigaster sulfurigaster* | SRR10100401 | 14.69 | 46.96 | 2 | 108 |
| *D. sulfurigaster neonasuta - pulaua* | SRR5059342 | 13.07 | 180.59 | 2 | 374 |
| *D. sulfurigaster neonasuta - pulaua* | SRR10100335 | 17.39 | 91.72 | 2 | 200 |
| *D. sulfurigaster neonasuta - D. sulfurigaster sulfurigaster* | SRR10100401 | 14.69 | 46.96 | 2 | 108 |
| *D. sulfurigaster neonasuta - D. sulfurigaster sulfurigaster* | SRR5059342 | 13.07 | 180.59 | 2 | 374 |
| *D. sulfurigaster sulfurigaster - D. pulaua* | SRR10100401 | 14.69 | 46.96 | 2 | 108 |
| *D. sulfurigaster sulfurigaster - D. pulaua* | SRR10100335 | 17.39 | 91.72 | 2 | 200 |
| *D. teissieri - D. erecta* | SRR5860615 | 12.82 | 7.06 | 2 | 26 |
| *D. teissieri - D. erecta* | SRR6425990 | 34.55 | 20.9 | 2 | 76 |
| *D. teissieri - D. orena* | SRR5860615 | 12.82 | 7.06 | 2 | 26 |
| *D. teissieri - D. orena* | SRR1977592 | 9.24 | 8.39 | 2 | 26 |
| *D. triauraria - D. rufa* | SRR13188922 | 16.05 | 12.22 | 2 | 40 |

| | | | | | |
|---|---|---|---|---|---|
| *D. triauraria - D. rufa* | SRR13070713 | 14.98 | 16.41 | 2 | 47 |
| *D. virilis - D. ezoana* | SRR20226095 | 22.75 | 10.2 | 2 | 43 |
| *D. virilis - D. ezoana* | SRR9426114 | 8.38 | 9.33 | 2 | 27 |
| *D. virilis - D. lacicola* | SRR9426114 | 8.68 | 8.47 | 2 | 25 |
| *D. virilis - D. lacicola* | SRR20226091 | 22 | 8.95 | 2 | 39 |
| *D. virilis - D. littoralis* | SRR9426114 | 9.94 | 8.52 | 2 | 26 |
| *D. virilis - D. littoralis* | SRR13070670 | 16.41 | 21.28 | 2 | 58 |
| *D. yakuba - D. erecta* | SRR9700025 | 16.48 | 7.73 | 2 | 31 |
| *D. yakuba - D. erecta* | SRR6425990 | 34.64 | 64.51 | 2 | 163 |
| *D. yakuba - D. orena* | SRR1977592 | 9.24 | 8.4 | 2 | 26 |
| *D. yakuba - D. orena* | SRR9700025 | 16.48 | 7.73 | 2 | 31 |
| *D. yakuba - D. santomea* | SRR9700025 | 16.48 | 7.73 | 2 | 31 |
| *D. yakuba - D. santomea* | SRR5860642 | 24.27 | 14.01 | 2 | 52 |
| *D. yakuba - D. teissieri* | SRR5860615 | 13 | 9.41 | 2 | 31 |
| *D. yakuba - D. teissieri* | SRR9700025 | 16.48 | 7.73 | 2 | 31 |

**Supplementary Table 4:** Species pairs and sample specific coverage information. Mean, standard deviation, minimum and maximum coverage are provided for each species in a pair-specific manner.

| Pair | Premating Isolation (Yukilevich 2014) | Postzygotic Isolation (Yukilevich 2014) | Nei's *D* *(Yukilevich 2014)* | Biogeographic classification (Yukilevich 2014) |
|---|---|---|---|---|
| *D. affinis - D. athabasca* | 0.775519466 | NA | 0.74 | Sympatry |
| *D. ambigua - D. obscura* | 0.993894994 | NA | 0.535 | Sympatry |
| *D. ambigua - D. persimilis* | NA | 1 | 1.66 | Allopatry |
| *D. ambigua - D. pseudoobscura* | NA | 1 | 1.66 | Allopatry |
| *D. americana - D. lacicola* | 1 | NA | 1.42 | Sympatry |
| *D. americana - D. littoralis* | 0.756484913 | NA | 1.21 | Allopatry |
| *D. americana - D. montana* | 0.992359932 | 1 | 1.48 | Sympatry |
| *D. americana - D. virilis* | 0.64428754 | 0 | 0.54 | Allopatry |
| *D. arizonae - D. mojavensis* | 0.766970618 | 0.375 | 0.279 | Allopatry |
| *D. auraria- D. rufa* | 0.987677141 | NA | 0.9245 | Sympatry |
| *D. bipectinata - D. malerkotliana* | 0.838798283 | 0.5 | 0.104 | Sympatry |
| *D. bipectinata - D. parabipectinata* | 0.55012987 | 0.5 | 0.145 | Sympatry |
| *D. bipectinata - D. pseudoananassae* | 0.840869565 | NA | 0.271 | Sympatry |
| *D. borealis - D. lacicola* | 1 | NA | 0.34 | Sympatry |
| *D. borealis - D. littoralis* | 1 | NA | 0.68 | Allopatry |
| *D. borealis - D. montana* | 0.994594595 | 0.5 | 0.21 | Sympatry |
| *D. erecta - D. orena* | 1 | NA | 1.03 | Sympatry |
| *D. flavomontana- D. montana* | 0.992857143 | 0.5 | 0.29 | Sympatry |
| *D. insularis - D. willistoni* | 0.923036649 | NA | 1.07 | Sympatry |
| *D. jambulina - D. punjabiensis* | 0.991326531 | NA | 0.944 | Sympatry |
| *D. kanekoi - D. ezoana* | 0.9945 | NA | 1.22 | Sympatry |
| *D. kanekoi - D. littoralis* | 0.5515 | NA | 1.22 | Allopatry |
| *D. kanekoi - D. lummei* | 0.722052705 | NA | 1.1 | Sympatry |
| *D. kanekoi - D. virilis* | 0.691660291 | NA | 1.1 | Sympatry |
| *D. kepulauna - D. albomicans* | 0.939064857 | 0 | 0.792 | Allopatry |
| *D. kikkawai- D. leontia* | 0.971875 | 0 | 0.244 | Sympatry |
| *D. kohkoa - D. albomicans* | 1 | 0.75 | 0.833 | Sympatry |
| *D. littoralis - D. ezoana* | 0.84125 | NA | 0.65 | Sympatry |

| | | | | |
|---|---|---|---|---|
| *D. littoralis - D. montana* | 0.784 | 0.75 | 0.66 | Sympatry |
| *D. loweii - D. pseudoobscura* | 0.98 | NA | 1.05 | Sympatry |
| *D. lummei - D. ezoana* | 1 | NA | 1.22 | Sympatry |
| *D. lummei - D. littoralis* | 0.629 | NA | 1.22 | Sympatry |
| *D. lummei - D. virilis* | 0.263 | 0 | 0.35 | Sympatry |
| *D. malerkotliana - D. parabipectinata* | 0.853214286 | 0.625 | 0.226 | Sympatry |
| *D. malerkotliana - D. pseudoananassae* | 0.956950673 | 0.95 | 0.282 | Sympatry |
| *D. mauritiana - D. erecta* | 0.984042553 | NA | 1.59 | Allopatry |
| *D. mauritiana - D. orena* | 0.9953125 | NA | 1.07 | Allopatry |
| *D. mauritiana - D. sechellia* | 0.920463693 | 0.5 | 0.32 | Allopatry |
| *D. mauritiana - D. teissieri* | 0.875842697 | NA | 1.24 | Allopatry |
| *D. mauritiana - D. yakuba* | 0.989247312 | NA | 0.88 | Allopatry |
| *D. melanogaster - D. erecta* | 1 | NA | 1.63 | Sympatry |
| *D. melanogaster - D. orena* | 1 | NA | 1.14 | Sympatry |
| *D. melanogaster - D. sechellia* | 0.619439164 | 1 | 0.62 | Allopatry |
| *D. melanogaster - D. simulans* | 0.826903313 | 1 | 0.55 | Sympatry |
| *D. melanogaster - D. teissieri* | 1 | NA | 1.01 | Sympatry |
| *D. melanogaster - D. yakuba* | 0.734042553 | NA | 0.94 | Sympatry |
| *D. miranda - D. persimilis* | 1 | 1 | 0.56 | Sympatry |
| *D. miranda - D. pseudoobscura* | 0.99537037 | 1 | 0.56 | Sympatry |
| *D. nasuta - D. albomicans* | 0.068400226 | 0 | 0.45 | Allopatry |
| *D. novamexicana - D. lacicola* | 1 | NA | 1.2 | Allopatry |
| *D. novamexicana - D. littoralis* | 0.788918206 | NA | 0.97 | Allopatry |
| *D. novamexicana - D. montana* | 1 | 0.75 | 1.22 | Sympatry |
| *D. novamexicana - D. virilis* | 0.507460036 | 0 | 0.5 | Allopatry |
| *D. obscura - D. tristis* | 0.989746683 | NA | 0.47 | Sympatry |

| | | | | |
|---|---|---|---|---|
| *D. parabipectinata - D. pseudoananassae* | 0.762171946 | NA | 0.366 | Sympatry |
| *D. persimilis - D. bifasciata* | 0.975 | NA | 1.95 | Allopatry |
| *D. persimilis - D. pseudoobscura* | 0.785042646 | 0.5 | 0.41 | Sympatry |
| *D. persimilis - D. subobscura* | 0.99137931 | NA | 1.66 | Allopatry |
| *D. pseudoobscura - D. bifasciata* | 0.941935484 | NA | 1.95 | Allopatry |
| *D. pseudoobscura - D. subobscura* | 0.962121212 | NA | 1.66 | Allopatry |
| *D. sechellia - D. erecta* | 1 | NA | 1.51 | Allopatry |
| *D. sechellia - D. orena* | 1 | NA | 1.27 | Allopatry |
| *D. sechellia - D. teissieri* | 0.995 | NA | 1.36 | Allopatry |
| *D. sechellia - D. yakuba* | 1 | NA | 1.27 | Allopatry |
| *D. simulans - D. erecta* | 0.997382199 | NA | 1.5 | Sympatry |
| *D. simulans - D. orena* | 0.953846154 | NA | 1.01 | Sympatry |
| *D. simulans - D. teissieri* | 0.961325967 | NA | 1.24 | Sympatry |
| *D. simulans - D. yakuba* | 0.958730159 | NA | 1 | Sympatry |
| *D. subobscura - D. ambigua* | 0.929569267 | NA | 1.44 | Sympatry |
| *D. subobscura - D. bifasciata* | 0.934782609 | NA | 1.33 | Allopatry |
| *D. subobscura - D. obscura* | 1 | NA | 1.25 | Sympatry |
| *D. subobscura - D. tristis* | 0.98273878 | NA | 1.1 | Sympatry |
| *D. sulfurigaster albostrigata - D. kepulauna* | 0.979381443 | 1 | 0.902 | Sympatry |
| *D. sulfurigaster albostrigata - D. pulaua* | 0.992078001 | 0 | 0.389 | Sympatry |
| *D. sulfurigaster albostrigata - D. sulfurigaster bilimbata* | 0.430365297 | 0.25 | 0.386 | Allopatry |
| *D. sulfurigaster albostrigata - D. sulfurigaster neonasuta* | 0.183673469 | 0 | 0.557 | Allopatry |
| *D. sulfurigaster albostrigata - D. sulfurigaster sulfurigaster* | 0.132816824 | 0 | 0.378 | Allopatry |
| *D. sulfurigaster bilimbata - D. pulaua* | 0.921381775 | 0 | 0.448 | Allopatry |

| | | | | |
|---|---|---|---|---|
| *D. sulfurigaster bilimbata - D. sulfurigaster neonasuta* | 0.595550485 | 0.5 | 0.491 | Allopatry |
| *D. sulfurigaster bilimbata - D. sulfurigaster sulfurigaster* | 0.097560976 | 0 | 0.555 | Allopatry |
| *D. sulfurigaster neonasuta - D. pulaua* | 0.909866017 | 0.5 | 0.705 | Allopatry |
| *D. sulfurigaster neonasuta - D. sulfurigaster sulfurigaster* | 0.415376106 | 0.5 | 0.6 | Allopatry |
| *D. sulfurigaster sulfurigaster - D. pulaua* | 0.785467128 | 0 | 0.561 | Allopatry |
| *D. teissieri- D. erecta* | 0.977011494 | NA | 1.54 | Sympatry |
| *D. teissieri - D. orena* | 1 | NA | 1.47 | Sympatry |
| *D. triauraria - D. rufa* | 0.963319303 | NA | 0.7335 | Sympatry |
| *D. virilis - D. ezoana* | 0.9645 | NA | 1.22 | Sympatry |
| *D. virilis - D. lacicola* | 0.799457995 | NA | 1.22 | Allopatry |
| *D. virilis - D. littoralis* | 0.682291667 | 0 | 1.22 | Allopatry |
| *D. yakuba - D. erecta* | 0.994505495 | NA | 1.4 | Sympatry |
| *D. yakuba - D. orena* | 1 | NA | 1.12 | Sympatry |
| *D. yakuba - D. santomea* | 0.842032011 | 0.5 | 0.3 | Sympatry |
| *D. yakuba - D. teissieri* | 0.988372093 | NA | 0.39 | Sympatry |

**Supplementary Table 5:** Information on reproductive isolation (pre- and postzygotic isolation), Nei's *D* based on allozymes and range overlap classifications for every pair considered in the analysis. These data are taken from Yukilevich (*3*, *4*).

| Pair | IM vs SI (Difference in logL -- No Recombination) | IM vs SI (Difference in logL -- With Recombination) |
|---|---|---|
| *D. affinis - D. athabasca* | -0.154304 | 9.4760575 |
| *D. ambigua - D. obscura* | -0.63247475 | 1.8216665 |
| *D. ambigua - D. persimilis* | -6.669142 | 0.22948725 |
| *D. ambigua - D. pseudoobscura* | -7.86996125 | 0.1155895 |
| *D. americana - D. montana* | 0.1722205 | 0.9437255 |
| *D. americana - D. virilis* | -0.00029675 | 1.1008325 |
| *D. americana - D. lacicola* | -5.675239 | 0.293779 |
| *D. americana - D. littoralis* | 0.061838 | 0.318727 |
| *D. arizonae - D. mojavensis* | -2.12492575 | 2.507734 |
| *D. auraria- D. rufa* | -5.53448175 | 1.8336325 |
| *D. bipectinata - D. malerkotliana* | -0.14979975 | 7.444685 |
| *D. bipectinata - D. parabipectinata* | 0.6888265 | 2.85252625 |
| *D. bipectinata - D. pseudoananassae* | 0.39919825 | 10.011253 |
| *D. borealis - D. lacicola* | -0.582845 | 3.011703 |
| *D. borealis - D. littoralis* | -6.40894375 | 0.89922525 |
| *D. borealis - D. montana* | -1.304686 | 1.46773075 |
| *D. erecta - D. orena* | 0.15642725 | 0.30473525 |
| *D. flavomontana- D. montana* | -1.47105625 | 0.28318675 |
| *D. insularis - D. willistoni* | -6.79094175 | 1.4804645 |
| *D. jambulina - D. punjabiensis* | 0.18039175 | 9.0430455 |
| *D. kanekoi - D. ezoana* | 0.491416 | 4.520206 |
| *D. kanekoi - D. littoralis* | -6.95620425 | 1.3461545 |
| *D. kanekoi - D. lummei* | 0.023294 | 4.623887 |
| *D. kanekoi - D. virilis* | -0.000216 | 0.25911775 |
| *D. kepulauna - D. albomicans* | -2.70380525 | 1.392413 |
| *D. kikkawai- D. leontia* | -5.1257895 | 0.885859 |
| *D. kohkoa - D. albomicans* | -0.05141025 | 8.10915425 |
| *D. littoralis - D. ezoana* | 0.4638655 | 7.7325425 |
| *D. littoralis - D. montana* | -4.93153525 | 0.9080145 |
| *D. loweii - D. pseudoobscura* | 0.1259705 | 1.475313 |
| *D. lummei - D. ezoana* | 0.19643625 | 5.336632 |
| *D. lummei - D. littoralis* | 0.08678375 | 8.1485735 |
| *D. lummei - D. virilis* | -0.0463755 | 7.52355625 |
| *D. malerkotliana - D. parabipectinata* | 0.13188 | 10.61952175 |
| *D. malerkotliana - D. pseudoananassae* | 0.29347375 | 3.239107 |

| | | |
|---|---|---|
| *D. mauritiana - D. erecta* | 0.3344945 | 0.608184 |
| *D. mauritiana - D. orena* | -0.00181775 | 0.22230825 |
| *D. mauritiana - D. sechellia* | -9.2088235 | 1.4877295 |
| *D. mauritiana - D. teissieri* | 0.0743675 | 1.60343475 |
| *D. mauritiana - D. yakuba* | -0.00088875 | 0.3945455 |
| *D. melanogaster - D. erecta* | 0.04074125 | 0.0600005 |
| *D. melanogaster - D. orena* | -0.000964 | -0.00286875 |
| *D. melanogaster - D. sechellia* | -0.0005805 | 0.4052805 |
| *D. melanogaster - D. simulans* | 0.7517305 | 0.68122625 |
| *D. melanogaster - D. teissieri* | 0.02389 | 1.208034 |
| *D. melanogaster - D. yakuba* | -1.30E-05 | 0.07559225 |
| *D. miranda - D. persimilis* | 0.4444185 | 1.53889375 |
| *D. miranda - D. pseudoobscura* | -0.0001285 | 1.50660875 |
| *D. nasuta - D. albomicans* | -1.63932875 | 1.9127715 |
| *D. novamexicana - D. lacicola* | -1.455242 | 4.48309925 |
| *D. novamexicana - D. littoralis* | -2.22167625 | 1.75211425 |
| *D. novamexicana - D. montana* | -5.5826495 | 0.73650975 |
| *D. novamexicana - D. virilis* | -4.97667175 | 0.451082 |
| *D. obscura - D. tristis* | 0.1502035 | 6.021555 |
| *D. parabipectinata - D. pseudoananassae* | 0.5475705 | 5.42126225 |
| *D. persimilis - D. bifasciata* | -5.97366225 | 3.8885565 |
| *D. persimilis - D. pseudoobscura* | 0.3772995 | 0.52908975 |
| *D. persimilis - D. subobscura* | -0.00042525 | 2.67E-05 |
| *D. pseudoobscura - D. bifasciata* | -1.89383775 | 0.03296825 |
| *D. pseudoobscura - D. subobscura* | 0.0205525 | 0.77406975 |
| *D. sechellia - D. erecta* | 0.106349 | 0.06987925 |
| *D. sechellia - D. orena* | -0.00011025 | 0.229013 |
| *D. sechellia - D. teissieri* | 0.01147 | 0.075828 |
| *D. sechellia - D. yakuba* | 0.016072 | 0.1990405 |
| *D. simulans - D. erecta* | -0.00114175 | 1.106863 |
| *D. simulans - D. orena* | -0.00078075 | 0.80872725 |
| *D. simulans - D. teissieri* | -0.00128975 | 1.1766165 |
| *D. simulans - D. yakuba* | -0.00290275 | 0.72976775 |
| *D. subobscura - D. ambigua* | 0.29490925 | 0.79274025 |
| *D. subobscura - D. bifasciata* | -7.03964625 | 0.2769065 |
| *D. subobscura - D. obscura* | 0.009463 | 0.18738925 |
| *D. subobscura - D. tristis* | -4.9943265 | 1.55378425 |
| *D. sulfurigaster albostrigata - D. kepulauna* | -5.10E-05 | 0.66901075 |
| *D. sulfurigaster albostrigata - D. pulaua* | -3.273116 | 1.75750475 |

| | | |
|---|---|---|
| *D. sulfurigaster albostrigata - D. sulfurigaster bilimbata* | -3.31889625 | 1.5914515 |
| *D. sulfurigaster albostrigata - D. sulfurigaster neonasuta* | -2.75170775 | 1.70006225 |
| *D. sulfurigaster albostrigata - D. sulfurigaster sulfurigaster* | -1.8848695 | 0.381667 |
| *D. sulfurigaster bilimbata - D. pulaua* | -1.76603975 | 1.33646125 |
| *D. sulfurigaster bilimbata - D. sulfurigaster neonasuta* | -2.7599 | 2.82219175 |
| *D. sulfurigaster bilimbata - D. sulfurigaster sulfurigaster* | -0.1085505 | 3.0562085 |
| *D. sulfurigaster neonasuta - D. pulaua* | -5.3976675 | 2.4274 |
| *D. sulfurigaster neonasuta - D. sulfurigaster sulfurigaster* | -0.73185425 | 0.7612595 |
| *D. sulfurigaster sulfurigaster - D. pulaua* | -1.39609025 | 1.44585225 |
| *D. teissieri - D. orena* | -0.00012325 | 0.31169375 |
| *D. teissieri- D. erecta* | 0.0013285 | 0.34871325 |
| *D. triauraria - D. rufa* | -1.820373 | 1.352234 |
| *D. virilis - D. ezoana* | 0.0004845 | 5.23917 |
| *D. virilis - D. lacicola* | -2.8273075 | 0.8447935 |
| *D. virilis - D. littoralis* | -7.478602 | 1.587921 |
| *D. yakuba - D. erecta* | -1.25E-06 | 0.9683835 |
| *D. yakuba - D. orena* | -0.00040575 | 0.36649375 |
| *D. yakuba - D. santomea* | -0.0001785 | 0.0026815 |
| *D. yakuba - D. teissieri* | -6.107981 | 1.13633825 |

**Supplementary Table 6:** Simulation inference. Difference in log likelihood for simulations without and with recombination. For each pair, we simulated a strict isolation with optimized parameters and with the same pair-specific block sizes as the observed dataset. We simulated blocks with and without realistic recombination within blocks and performed model fitting to determine the potential effect of intra-block recombination on our inference. We compare the support for an IM model over the SI model. Values shown are the differences in logL between SI and IM model for each pair in simulations without and with recombination under a strict isolation model.
